# β1 integrin signaling governs necroptosis via the chromatin remodeling factor CHD4

**DOI:** 10.1101/2023.04.14.536920

**Authors:** Zhiqi Sun, Filippo M. Cernilogar, Helena Horvatic, Assa Yeroslaviz, Zeinab Abdullah, Gunnar Schotta, Veit Hornung

## Abstract

Fibrosis, characterized by sustained activation of myofibroblasts and excessive extracellular matrix (ECM) deposition, is known to be associated with chronic inflammation. RIPK3, the central kinase of necroptosis signaling, is upregulated in fibrosis and contributes to the TNF-mediated inflammation. In bile duct ligation-induced liver fibrosis, we found that myofibroblasts are the major cell type expressing RIPK3. Genetic ablation of β1 integrins, the major profibrotic ECM receptors in fibroblasts, not only abolished ECM fibrillogenesis but also blunted RIPK3 expression via an epigenetic mechanism mediated by the chromatin remodeling factor CHD4. While the function of CHD4 has been conventionally linked to NuRD and ChAHP complexes, we found that CHD4 potently repressed a set of genes, including *Ripk3*, with high locus specificity but independent of either the NuRD or ChAHP complex. Thus, our data uncover that β1 integrin intrinsically links fibrotic signaling to RIPK3-driven inflammation via a novel mode of action of CHD4.

## Introduction

Tissue fibrosis is initiated by a prolonged wound healing response during chronic tissue injury and inflammation. It is characterized by sustained activation of contractile myofibroblasts that mediate excessive deposition of extracellular matrix (ECM) such as collagen and fibronectin and tissue stiffening *(1)*. As the major receptors mediating cell adhesion to the ECM, integrins in myofibroblasts play a pivotal role in fibrosis *(2, 3)*. 24 heterodimeric integrin receptors are expressed in vertebrates. Among these, a group of β1-containing integrin receptors bind collagens (α1β1, α2β1, α10β1, α11β1) and fibronectin (α5β1), mediate ECM fibrillogenesis, promote cellular contractility, and instruct a profibrotic gene expression program through the MKL1-SRF and YAP/TAZ transcription factor networks *(4–7)*. As such, they can promote an aberrant mechano-transduction signaling state in fibrosis. In fact, blocking β1 integrin prevents ECM fibrillogenesis, reduces cellular contractility, and blunts profibrotic gene expression *(5, 7, 8)*. In contrast to β1 integrin, αVβ3 integrin in fibroblasts binds fibronectin but not collagen, cannot mediate fibrillogenesis, but can synergize with β1 integrins to promote cellular contractility *(4)*.

Myofibroblast activation during tissue injury and wound healing is driven by a myriad of profibrotic and proinflammatory cytokines most notably transforming growth factor-β (TGFβ), interleukin-1β (IL-1β) and tumor-necrosis factor α (TNFα). Binding of TNFα to TNFR1 triggers receptor-interacting protein kinase 1 (RIPK1)-dependent NF-κB signaling that promotes the expression of anti-apoptotic proteins such as BCL2L1, c-FLIP and cIAP. Prolonged TNFR1 activation triggers activation of caspase-8 and the immune-silent cell death of apoptosis. Insufficient caspase-8 activity, however, results in the formation of a large amyloid-like signaling complex via RHIM-motif-mediated oligomerization of RIPK1 and RIPK3 *(9–11)*. Subsequently, activated RIPK3 phosphorylates pseudokinase mixed lineage kinase domain-like protein (MLKL), which then oligomerizes at the plasma membrane to induce membrane rupture and a lytic form of cell death known as necroptosis *(12–14)*. During necroptosis, dying cells unleash damage-associated molecular patterns (DAMPs) and alarmins such as IL-1 family cytokines, and TNFα *(15–17)*. Furthermore, RIPK3 can cooperate with RIPK1 to promote a large wave of cytokine expression, thereby instigating a self-amplifying inflammatory signaling loop that impacts on the cell microenvironment beyond cell death *(18)*. Blocking TNFα signaling has been proven beneficial in fibrosis in multiple organs including liver, lung, and kidney, and in fibrosis associated with Duchenne muscular dystrophy (DMD) and Dupuytren’s disease *(19–21)*.

Interestingly, RIPK3 upregulation is associated with disease progression in large cohort of human non-alcoholic steatohepatitis (NASH) patients *(22)*. RIPK3 upregulation has also been reported in lung fibrosis *(23)*, kidney fibrosis *(24)* and in Duchenne muscular dystrophy (DMD), which is associated with muscle fibrosis *(21, 25)*, suggesting a universal mechanism driving RIPK3 expression in fibrosis. The cell types that express RIPK3 within fibrotic tissues are controversial. While RIPK3 upregulation has been attributed to fibroblasts in renal fibrosis *(24)*, RIPK3 remains silenced by promoter methylation in hepatocytes in non-alcoholic steatohepatitis (NASH) *(26)*, suggesting that non-hepatic cells may contribute to RIPK3 upregulation in liver fibrosis. Nevertheless, genetic ablation of RIPK3 alleviates unilateral ureteral obstruction–induced or adenine diet-induced renal fibrosis *(24)*, methionine-and choline-deficient diet-induced steatohepatitis *(27)* and DMD-associated muscle fibrosis *(21)*.

How RIPK3 expression is regulated in fibrosis remains poorly understood. While the RIPK3 locus is silenced by DNMT1-mediated promoter methylation in *in vitro* cultured cancer cell lines *(28)*, the CHD4-containing NuRD (Nucleosome Remodeling Deacetylase) complex has been reported to repress RIPK3 expression in endothelial cells and muscle stem cells *(25, 29)*. The NuRD complex is composed of at least 7 subunits, each with at least 2 paralogs, including Class-II Chromodomain helicase DNA-binding (CHD) proteins CHD3/4/5, GATAD2A/B, CDK2AP1/2, MBD2/3, MTA1/2/3, RBBP4/7 and HDAC1/2 *(30)*. By CHD-dependent nucleosome sliding into a dense array and HDAC1/2-mediated erasure of active transcription marks such as H3K27Ac, NuRD generally functions as a transcriptional repressor. Class-II CHDs also form another, recently described complex with activity dependent neuroprotector homeobox (ADNP) and heterochromatin protein 1 (HP1), known as the ChAHP complex *(31, 32)*. Together, NuRD and ChAHP constitute most of the high-affinity interactors of Class-II CHDs *(32, 33)* and are thought to account for the majority of their functions in stem cell maintenance, cell lineage specification, Kaposi sarcoma-associated herpesvirus (KSHV) latency, hemoglobin switch, and DNA damage repair *(34–39)*. On the other hand, instead of being a core component of the NuRD complex, CHD4 only peripherally associates with the NuRD complex *(40, 41)* and has nucleosome-remodeling activity by itself *(42)*, suggesting a possible NuRD-and ChAHP-independent function.

Here, we investigated the cell autonomous regulation between fibrotic signaling and RIPK3-mediated TNF signaling in fibroblasts. We found strong RIPK3 expression in both myofibroblasts and macrophages in liver fibrosis induced by bile duct ligation. Genetic ablation of the profibrotic β1 integrin receptor in fibroblasts not only abolished the fibrotic signature but also reduced RIPK3 expression through CHD4-mediated repression at the proximal enhancer element within the RIPK3 locus. Surprisingly, despite its high locus specificity, CHD4-mediated repression of RIPK3 operated independently of the NuRD or ChAHP complex, arguing for the existence of additional CHD4-containing complexes that regulate locus-specific gene expression.

## Results

### β1 integrin links fibrotic signalling with RIPK3 expression in fibroblasts

Numerous reports of RIPK3 upregulation in liver fibrosis and the lack of necroptosis in hepatocytes prompted us to define the spatial expression of RIPK3 in a bile duct ligation (BDL)-induced liver fibrosis model. Whereas RIPK3 was sparsely detected in the healthy liver, most likely in Kupffer cells, RIPK3 expression was significantly increased in fibrotic liver tissue, mostly within fibrotic scars where SMA-positive myofibroblasts and infiltrating CD11b-positive myeloid cells were tightly intercalated, but also slightly in the surrounding hepatocytes (Fig. S1A). While RIPK3 is known to be expressed in macrophages, we observed comparably high RIPK3 expression in myofibroblasts (Fig. S1B). Immunostaining with another RIPK3 antibody confirmed that RIPK3 was upregulated in BDL-induced liver fibrosis colocalizing with the mesenchyme marker Vimentin (Fig. S1C & D). Compared with healthy liver, the mean fluorescence intensity of RIPK3 immunolabeling was increased 3-fold in the fibrotic area in parallel with Vimentin signals, and moderately increased by 50% in the surrounding area (Fig. S1E & F).

Myofibroblasts deposit excessive ECM and mediate tissue stiffening through integrin receptors. To test whether fibrotic signaling is intrinsically linked to RIPK3 signaling and necroptosis in fibroblasts, we used a set of well-established integrin pan-KO fibroblast cell lines in which fibrillogenic β1 integrin and non-fibrillogenic αV integrin are reconstituted individually (hereafter referred to as pKO-β1 and pKO-αV, respectively) or in combination (pKO-αV/β1) *(4)*. Indeed, immunostaining of confluent culture of these cells for collagen type 3 confirmed that only pKO-β1 and pKO-αV/β1, but not pKO-αV cells assembled elaborated collagen-3 ECM fibers (Fig. 1A). Intriguingly, combined treatment of TNFα and pan-caspase inhibitor zVAD triggered massive necroptosis in pKO-β1 cells as revealed by CytoxGreen labeling, whereas the non-fibrotic pKO-αV cells were completely resistant to necroptosis (Fig. 1A). Fibroblasts expressing both αV and β1 integrins showed an intermediate phenotype with moderate necroptosis (Fig. 1A). Live cell imaging with CytoxGreen revealed a hypersensitive kinetics of necroptosis in pKO-β1 cells that started 3 hours after TNFα/zVAD treatment (Fig. 1B). In contrast, minimal lytic cell death was observed in pKO-αV cells even after a period of 10 hours (Fig. 1B). An LDH release assay further confirmed the complete lack of necroptosis in pKO-αV cells (Fig. 1C). The differential necroptosis response is not caused by altered RIPK1 signaling, as comparable RIPK1-dependent induction of *Cxcl10* and *Cxcl1* mRNAs were observed in the three cell lines (Fig. S1G). Instead, the degree of necroptosis correlated with MLKL phosphorylation at Ser345 by RIPK3 (Fig. 1D). Interestingly, whereas high and intermediate levels of RIPK3 protein were expressed in pKO-β1 and pKO-αV/β1 cell respectively, no RIPK3 protein expression could be detected in pKO-αV cells (Fig. 1D), suggesting a positive correlation between β1 integrin and RIPK3 expression in these cells.

**Figure 1.**
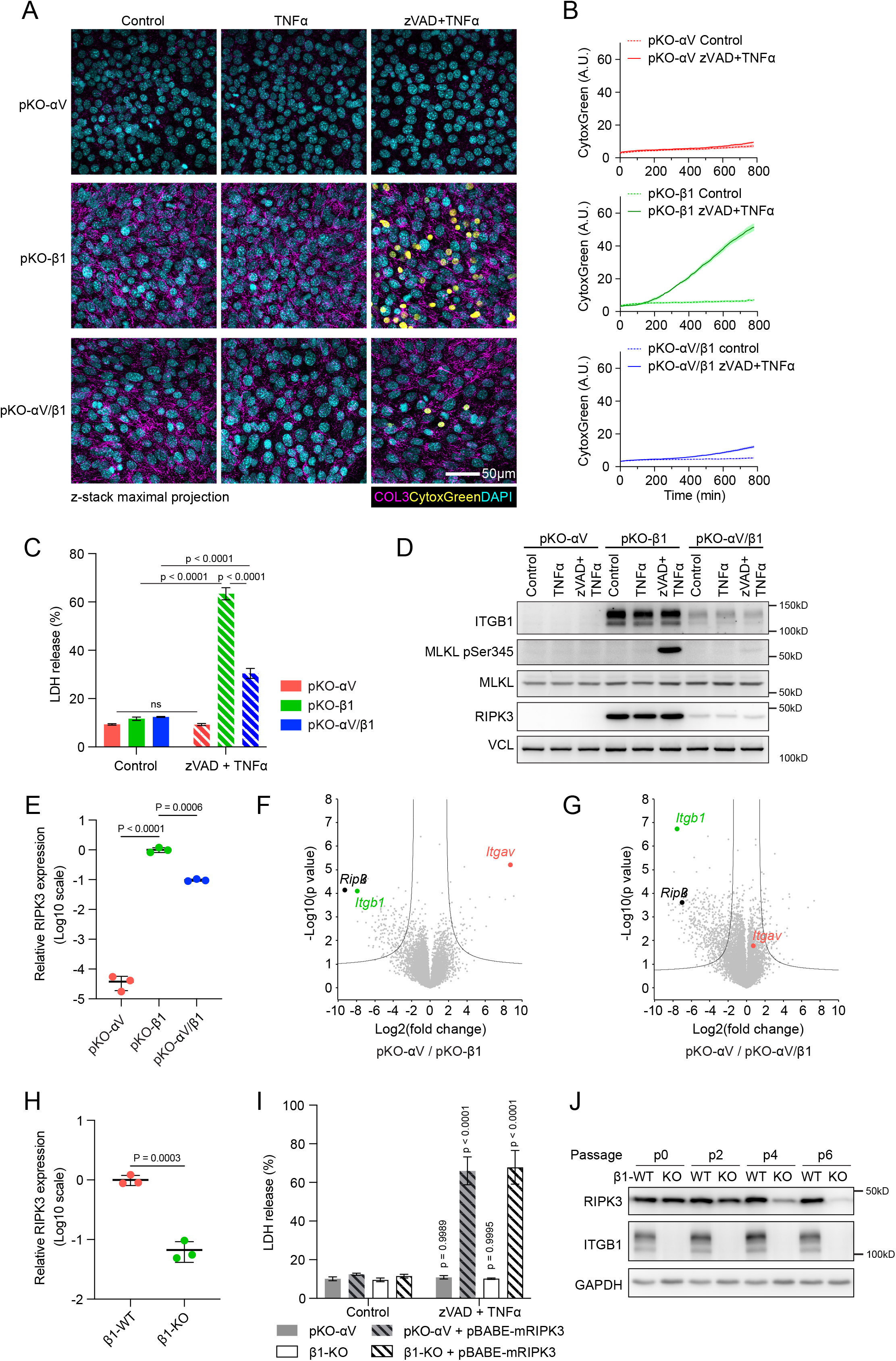
β1 integrin links fibrotic signalling to RIPK3 expression in fibroblasts. **(A)** pKO-αV, pKO-αV/β1 and pKO-β1 cells were seeded on FN-coated (10 μg/ml) coverslips for 24 hours at confluence and treated with vehicle control, 50 ng/ml TNFα or 40 µM zVAD and 50 ng/ml TNFα in combination for a further 8 hours. Cells were briefly stained with CytoxGreen (pseudocoloured in yellow) for 1 hour before fixation with paraformaldehyde and immunostained for COL3 (magenta) and DAPI (cyan). Maximum intensity projection of z-stack image series. Scale bar, 50 μm. **(B)** pKO-αV, pKO-αV/β1 and pKO-β1 cells were seeded on FN-coated 24-well plates for 24h at confluence and treated with vehicle control or 40 µM zVAD and 50 ng/ml TNFα in combination in the presence of CytoxGreen. Time lapse images were taken every 5 minutes for 24 hours. The time course of CytoxGreen signal intensity is shown as a line graph of the mean ± s.e.m. of three replicates. **(C)** pKO-αV, pKO-αV/β1 and pKO-β1 cells were seeded on FN-coated 24-well plates for 24 hours at confluence and treated with vehicle control or 40 µM zVAD and 50 ng/ml TNFα in combination for 18 hours. Normalized LDH release values are depicted as mean ± s.d. (n = 3); p value calculated by two-way ANOVA with Tukey correction for multiple testing. **(D)** Cells were treated as in **(A)** and protein expression of ITGB1, MLKL, MLKL-pSer345 was analysed by Western blot. VCL was used as loading control. Data are representative of 2 independent experiments. **(E)** qPCR analysis of *Ripk3* transcript in pKO-αV, pKO-αV/β1 and pKO-β1 cells. Data were normalised to *Gapdh* and are depicted as mean ± s.d. (n = 3); p values were calculated on log-transformed data by one-way ANOVA with Tukey correction for multiple testing. **(F)** & **(G)** Volcano plot comparison of DEseq2 count of RNA-seq data between pKO-αV and pKO-β1 cells **(F)** and between pKO-αV and pKO-αV/β1 cells **(G)**. **(H)** qPCR analysis of *Ripk3* transcript in β1-WT and β1-KO fibroblasts. Data were normalised to *Gapdh* and are depicted as mean ± s.d. (n = 3); p values were calculated on log-transformed data by an unpaired student’s t-test. **(I)** pKO-αV, β1-KO and corresponding cell line reconstituted with mouse RIPK3 were treated as in **(C)** and LDH release was analysed and is depicted as mean ± s.d. (n = 3); p values were calculated by two-way ANOVA with Šídák’s correction for multiple testing (for every cell line, a comparison of unstimulated and stimulated cells was conducted). **(J)** β1-integrin floxed fibroblasts were infected with adenovirus expressing Cre recombinase for 2 days. Freshly sorted β1-KO cells were designated as passage 0 and passaged every 2 days. Total cell lysate from both β1-WT and β1-KO cells was collected between passage 0 and passage 6 and protein expression of ITGB1, RIPK3 was analysed by Western blot. GAPDH was used as a loading control. Data are representative of 2 independent experiments.

qPCR analysis revealed that while high and intermediate levels of *Ripk3* mRNA were detected in pKO-β1 and pKO-αV/β1 cells, *Ripk3* mRNA expression was markedly repressed in pKO-αV cells (Fig. 1E). Strikingly, unbiased RNA-seq analysis of the three integrin-reconstituted cell lines revealed that *Ripk3* mRNA was the most significantly upregulated transcript in β1 integrin-expressing cells (Supplementary Table 3, Fig. 1F & G). To further confirm these results, we used an SV40 large T antigen-immortalized fibroblast cell line derived from β1 integrin floxed mice (hereafter referred to as β1-WT) *(43)*. Cre recombinase-mediated deletion of endogenous β1 integrin (β1-KO) led to significant reduction of *Ripk3* mRNA expression in comparison with the parental cells (Fig. 1H). Like pKO-αV cells, β1-KO fibroblasts were entirely resistant to TNFα-induced necroptosis which could be rescued by lentivirus-mediated reconstitution of RIPK3, indicating that RIPK3 is the only missing component for necroptosis in β1 integrin-deficient cells (Fig. 1I). Notably, RIPK3 reconstitution in pKO-αV and β1-KO cells did not trigger basal cytotoxicity (Fig. 1I), indicating that the loss of RIPK3 expression in β1-deficient cells was not due to RIPK3-induced cell death. To further understand how β1 integrin deficiency leads to diminished RIPK3 expression, we infected an early passage of β1 integrin floxed fibroblasts with adenovirus expressing Cre recombinase and FACS-sorted β1-KO cells by surface labeling 2 days after Cre expression. Sorted cells (designated as P0) were passaged every 2 days thereafter. Whereas RIPK3 expression was maintained after β1 integrin deletion in P0 cells, reduction of RIPK3 became discernible at passage 2 (P2), which was succeeded by slow but progressive loss of RIPK3 in the following passages (Fig. 1J). In conclusion, our data indicate that RIPK3 expression and thus sensitivity to necroptosis-inducing stimuli is profoundly affected by β1 integrin signaling. Furthermore, our data also suggest that this phenomenon is exerted via a persistent epigenetic mechanism.

### CHD4 but not DNMTs represses RIPK3 expression in β1 integrin-deficient cells

DNMT-mediated promoter methylation was shown to epigenetically suppress *Ripk3* expression in various *in vitro* cultured cancer cell lines. Although inhibition of DNMT activity by 5-Aza-2’-deoxycytidine (5’AZA) moderately increased *Ripk3* mRNA expression, this treatment did not normalize *Ripk3* mRNA levels in integrin-reconstituted cells (Fig. S2A). Similarly, DNMT inhibition failed to restore *Ripk3* mRNA expression of β1-KO cells to the level of the parental wildtype cells (Fig. S2B). The CHD4/NuRD complex was recently shown to suppress RIPK3 in muscle stem cells and endothelial cells *(25, 29)*. Notably, knockdown of CHD4 by three different shRNAs significantly increased RIPK3 expression and fully normalized RIPK3 expression between wildtype and β1-KO cells, both at the protein level (Fig. 2A & B) and at the mRNA level (Fig. 2C). Moreover, live cell imaging revealed that CHD4 depletion rendered necroptosis-resistant β1-KO cells hypersensitive to TNFα/zVAD-induced necroptosis (Fig. 2D). Equal protein expression (Fig. 2A) and nuclear localization (Fig. S2C) of CHD4 were observed in β1-WT and β1-KO cells, suggesting that β1 integrin may regulate CHD4 activity rather than its expression or localization. We further tested whether CHD4-mediated RIPK3 repression is a conserved mechanism in other cell types. Transient knockdown of CHD4 by siRNAs significantly increased RIPK3 expression in β1-KO fibroblasts, B16-F10 melanoma cells, Panc02 PDAC cells and MOVAS smooth muscle cell. However, cells with higher basal RIPK3 expression tended to be less responsive to CHD4 depletion in terms of RIPK3 upregulation. Notably, L929 cells expressed the highest levels of RIPK3 among the cell lines tested, at levels comparable to that in *Chd4* KO fibroblasts, which could not be further increased by CHD4 knockdown (Fig. S2D & E).

**Figure 2.**
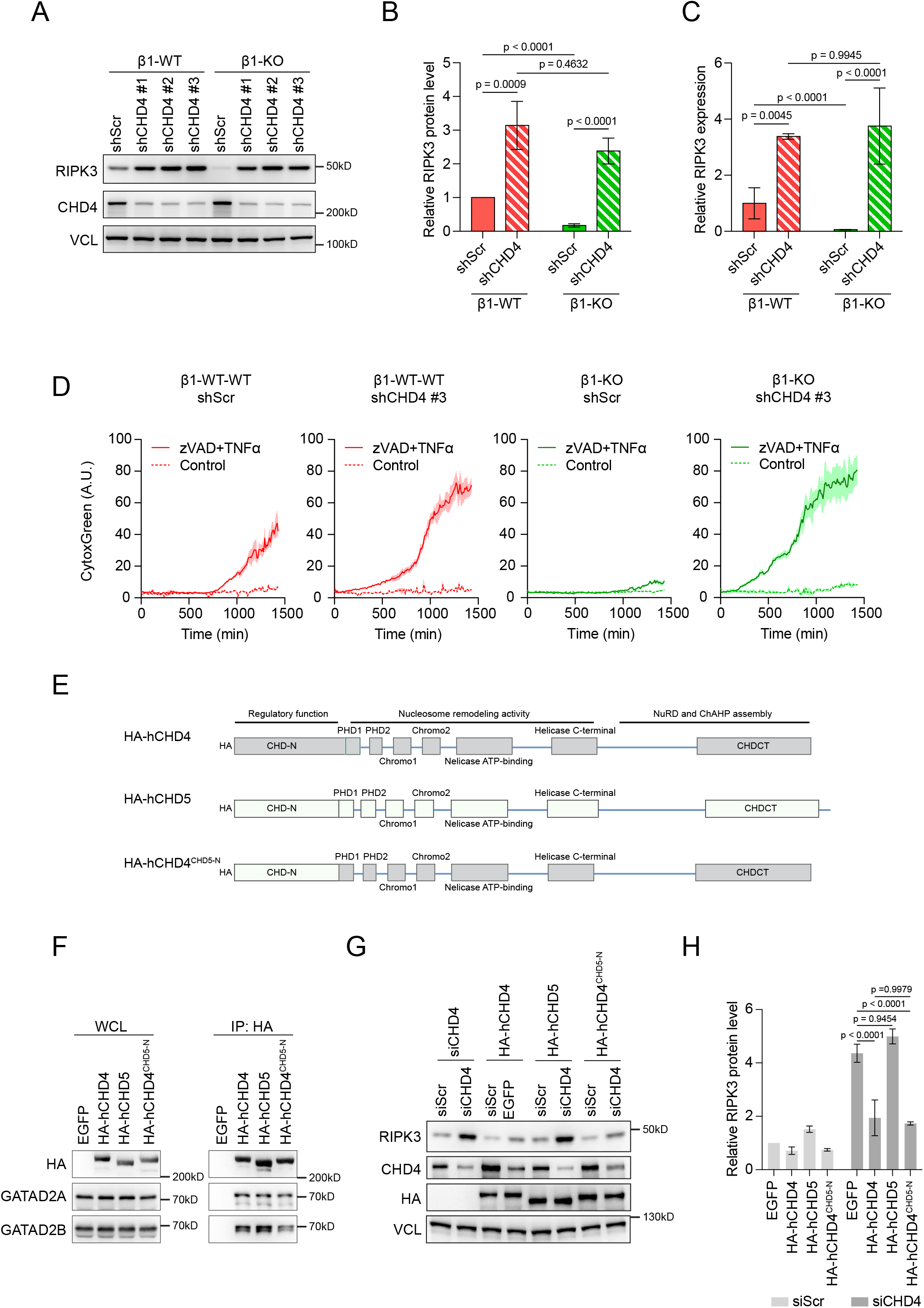
CHD4 represses RIPK3 expression in β1-integrin-deficient cells. **(A)** β1-WT and β1-KO fibroblasts are stably transduced with lentivirus expressing scramble control shRNA (shScr) or three different shRNA targeting mouse CHD4 (shCHD4 #1,2,3). Protein expression of RIPK3 and CHD4 was analysed by Western blot. VCL was used as a loading control. Data are representative of 3 independent experiments. **(B)** RIPK3 protein expression level from cells in **(A)** was quantified by densitometry and normalised to that in β1-WT expressing shScr and are depicted as mean ± s.d. (n = 3); p values were calculated on log-transformed data by a two-way ANOVA with Tukey correction for multiple testing. **(C)** qPCR analysis of *Ripk3* transcript expression in β1-WT and β1-KO fibroblasts expressing control scramble shRNA or shRNA against CHD4. Data are normalized to the mean value of the β1-WT / scramble shRNA controls and depicted as mean ± s.d. (n = 3); p values were calculated on log-transformed data by a two-way ANOVA with a Tukey correction for multiple testing. **(D)** β1-WT and β1-KO fibroblasts expressing control scramble shRNA or shCHD4 #3 were seeded on FN-coated 24-well plates for 24 hours at confluence and treated with vehicle control or 40 µM zVAD and 50 ng/ml TNFα in combination in the presence of CytoxGreen. Time lapse images were taken every 5 minutes for 24 hours. The time course of CytoxGreen signal intensity is shown as a line graph of the mean ± s.e.m. of three replicates. **(E)** Domain architecture of human CHD4, CHD5 and the design of a chimeric CHD4 with its N-terminal regulatory region replaced by that of hCHD5 (hCHD4^hCHD5-N^). The nucleosome remodelling activity is located in the central part containing tandem PHD, chromo domains and helicase domain. Assembly interfaces for NuRD and ChAHP complexes are located in the C-terminal region. **(F)** β1-KO fibroblasts were stably transduced by the piggy bac transposon to express EGFP control, N-terminal HA-tagged hCHD4, hCHD5 and hCHD4^hCHD5-N^ as shown in **(E)**. HA-tagged CHD proteins were enriched by anti-HA beads and co-immunoprecipitated GATAD2A and GATAD2B were analysed by Western blot. Data are representative of 2 independent experiments. **(G & H)** Non-targeting control siRNA (siScr) or siRNA against mouse CHD4 (siCHD4) were transiently transfected for 72h into the cell lines established in **(F)**. Protein expression of CHD4 and HA-tagged CHDs as well as RIPK3 was analysed by Western blot using VCL as loading control **(G)**. Data are representative of 3 independent experiments. **(H)** Relative RIPK3 expression levels were quantified by densitometric analysis and are depicted as mean ± s.d. (n = 3); p values were calculated on log-transformed data by a two-way ANOVA with a Tukey correction for multiple testing.

Class-II CHDs including CHD3, CHD4 and CHD5 share a similar domain architecture, high sequence homology and they can all assemble NuRD and ChAHP complexes (Fig. 2E). Whereas the nucleosome binding and remodeling activity resides in the central part of the protein containing tandem PHD domains and Chromo domains followed by ATP-dependent helicase domain *(42)*, the assembly interface for NuRD and ChAHP resides in their C-terminal region. The less conserved N-terminal domain binds RNAs and plays a regulatory role *(38, 44)*. To test whether other class-II CHDs also regulate RIPK3 expression, EGFP control, HA-tagged human CHD4, CHD5 or a chimeric CHD4 containing N-terminal regulatory domain of CHD5 (CHD4^CHD5-N^) were stably expressed in β1 integrin KO fibroblasts (Fig. 2E). Immuno-precipitation of HA-tagged proteins confirmed that all CHD constructs were equally expressed and assembled into a GATAD2A/B-containing NuRD complex (Fig. 2F). Interestingly, while transient knockdown of endogenous CHD4 by siRNAs targeting murine *Chd4* mRNA increased RIPK3 expression in cells expressing EGFP control and hCHD5, stable expression of hCHD4 or hCHD4^CHD5-N^ prevented RIPK3 upregulation upon knockdown of endogenous CHD4 (Fig. 2G & H). Taken together, these results indicate that loss of β1 integrin signaling results in CHD4-mediated repression of *Ripk3* transcription, a phenomenon that is well conserved across different cell types. Furthermore, our data showed that CHD4 represses RIPK3 expression in a non-redundant manner within the class II CHD family proteins and that this function is conserved between mouse and human.

### CHD4 controls RIPK3 expression independent of NuRD or ChAHP complexes

The non-redundant function of CHD4 prompted us to test the involvement of NuRD and ChAHP, two major CHD4-associated chromatin remodeling complexes. Recent biochemical analysis revealed that CHD4 is linked to the remainder of the NuRD complex through GATAD2A/B with a possible contribution from CDK2AP1/2 *(30, 32)*. We therefore decoupled CHD4 from NuRD complex by stable knockdown of GATDA2A/B and CDK2AP1/2 proteins (Fig. 3A). The knockdown efficiency and protein expression levels of all NuRD subunits were profiled by label-free quantification (LFQ) through mass spectrometry (Supplementary Table 4 & 5). For the ease of comparison, sums of LFQ intensity of paralogs are summarized in Figure 3B. CHD4 knockdown led to >80% reduction of CHD4 and CDK2AP1/2 proteins, and approximately 50% reduction of other NuRD subunits including GATAD2A/B, MBD2/3, MTA1/2/3, and HDAC1/2. GATAD2A/B double knockdown resulted in >80% reduction of GATAD2A/B and CDK2AP1/2 proteins and approximately 50% reduction of other NuRD components including CHD3 and CHD4. CDK2AP1/2 knockdown predominantly affected protein level of CHD3 (around 80% reduction) with only mild effect on other NuRD subunits. Notably, ADNP levels were reduced by 50% upon CHD4 knockdown but remained intact upon GATAD2A/B or CDK2AP1/2 depletion. Paralog-specific effects were also documented (Fig. S3A). For instance, CHD4 depletion preferentially reduced GATAD2A but not GATAD2B; and MTA2 but not MTA1 was sensitive to GATAD2A/B or CHD4 depletion. Taken together, these data confirm mutual stabilization between NuRD subunits and independence between NuRD and ChAHP complexes. Moreover, depletion of the majority of CHD4 and GATAD2A/B by their respective shRNAs caused comparable disruption of the rest of the NuRD complex.

**Figure 3.**
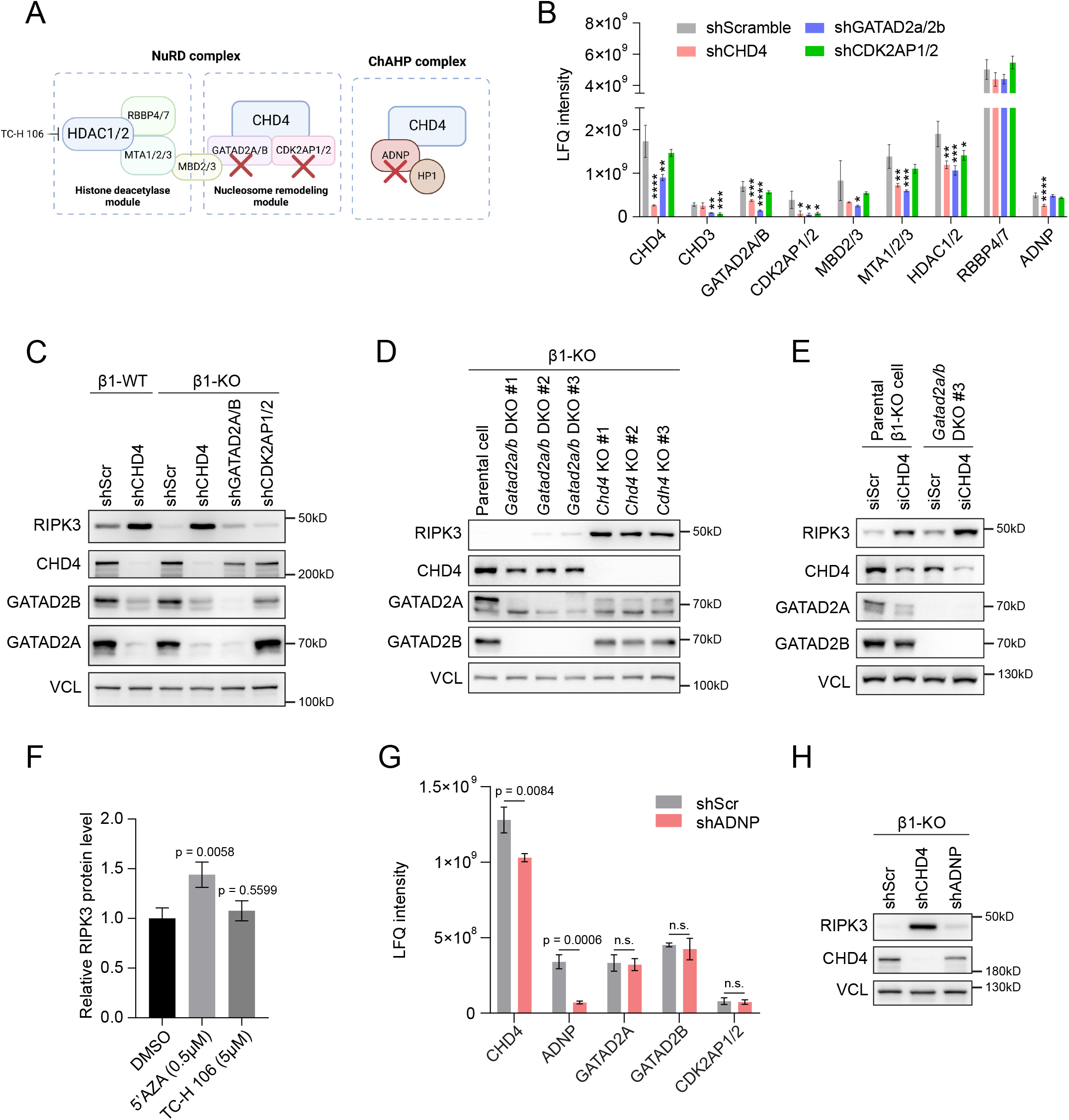
CHD4 controls RIPK3 expression independently of NuRD or ChAHP complexes. **(A)** Diagram of the modular assembly of the NuRD and ChAHP complexes. CHD4 is linked to the rest of the NuRD complex by GATAD2A/B with possible contribution of CDK2AP1/2. ADNP links CHD4 to HP1 in the ChAHP complex. GATAD2A/B and CDK2AP1/2 were targeted to uncouple CHD4 from the rest of the NuRD complex, while ADNP was targeted to disrupt the ChAHP complex. TC-H 106 was used to inhibit HDAC1/2 activity. **(B)** Sum of LFQ intensity of paralogous subunits of NuRD and ChAHP in β1-KO cells stably expressing control scramble shRNA, shRNAs targeting GATAD2A/B or CDK2AP1/2. Data are depicted as mean ± s.d. (n = 3); p values were calculated by one-way ANOVA with a Dunnett correction for multiple testing (for each protein or paralogous group, a comparison between the shRNA control and the other knockdown conditions was conducted). *, p < 0.05; **, p < 0.01; ***, p < 0.001; ****, p < 0.0001. **(C)** Immunoblot analysis of RIPK3, CHD4, GATAD2A, GATAD2B using VCL as loading control in β1-WT and β1-KO cells expressing control scramble shRNA or shRNA targeting CHD4, GATAD2A/B or CDK2AP1/2. Data are representative of 2 independent experiments. **(D)** Immunoblot analysis of RIPK3, CHD4, GATAD2A, GATAD2B in parental β1-KO cells, three *Chd4* KO clones and three *Gatad2a/b* DKO clones using VCL as loading control. Data are representative of 2 independent experiments. **(E)** Parental β1-KO cells and one *Gatad2a/b* DKO clone were transiently transfected with non-targeting control siRNA or siRNA against mouse CHD4 for 72 hours. Protein expression of CHD4, GATAD2A, GATAD2B and RIPK3 was analysed by immunoblot using VCL as a loading control. **(F)** β1-KO cells were treated with DMSO control, 0.5 µM 5-Aza-2’-desoxycytidin (5’AZA) or 5 µM TC-H 106 for 72 hours. The protein expression level of RIPK3 was analysed by immunoblot and densitometry analysis. p values were calculated on log-transformed data by a one-way ANOVA with Dunnett correction for multiple testing (a comparison of the DMSO control with the other conditions was conducted). **(G)** LFQ intensity of NuRD and ChAHP subunits in β1-KO cells stably expressing control scramble shRNA or shRNA against ADNP. Data are presented as mean ± s.d. (n = 3); p values were calculated by a Student t-test. **(H)** Immunoblot analysis of RIPK3, CHD4 using VCL as loading control in β1-KO cells expressing control scramble shRNA, shRNA against CHD4 or ADNP.

Western blot of CHD4, GATAD2A and GATAD2B confirmed the results of mass spectrometry (Fig. 3C). Interestingly, while CHD4 knockdown restored RIPK3 expression in β1 integrin KO cells, knockdown of GATAD2A/B or CDK2AP1/2 only slightly increased RIPK3 expression likely due to the partial destabilization of CHD4 (Fig. 3C). To further confirm this result, we generated *Chd4* KO and *Gatad2a/b* DKO in β1-KO cells. Consistently, *Chd4* KO or *Gatad2a/b* DKO led to a partial destabilization of each other (Fig. 3D). In contrast to the drastic upregulation of RIPK3 in *Chd4* KO clones, only slight increase of RIPK3 could be observed in *Gatad2a/b* DKO clones (Fig. 3D). Importantly, transient knockdown of CHD4 in *Gatad2a/b* DKO cells could strongly increase RIPK3 expression, with an effect size comparable to that of CHD4 knockdown in the parental β1-KO cells (Fig. 3E). Finally, RIPK3 expression was not altered by 3 days treatment with the class-I HDAC inhibitor, TC-H 106 (Fig. 3F). These data demonstrate that CHD4 suppresses RIPK3 expression independently of the NuRD complex or its associated HDAC1/2 activity.

We next disrupted the ChAHP complex by stable knockdown of ADNP (Fig. 3A). ADNP2 was not targeted since we did not detect its peptide in our cells in the previous MS analysis. ADNP knockdown resulted in 80% reduction of ADNP protein and mild reduction of CHD4 by 20%, without affecting protein levels of other NuRD subunits (Fig. 3G, Supplementary Table 6), confirming that the ChAHP and NuRD complex are independent from each other. RIPK3, however, was only marginally increased in ADNP-depleted cells likely due to the mild reduction of CHD4 (Fig. 3H). In conclusion, our data indicated that CHD4-mediated suppression of RIPK3 expression is independent of the NuRD or ChAHP complex.

### CHD4 exhibits high locus specificity independent of NuRD and ChAHP

Since CHD4 binds to the nucleosome and DNA without sequence preference, we investigated whether CHD4 could exhibit any locus-specificity independent of the NuRD or ChAHP complex. Whole proteome comparison between β1 integrin KO cells expressing a scramble shRNA or a CHD4 shRNA revealed that the LFQ intensities of 1,113 proteins out of 6,498 quantified proteins were significantly altered (FDR < 0.05, S0 = 0.1, Supplementary Table 4 & 5). Notably, besides RIPK3, a group of proteins related to the extracellular matrix, cytoskeleton and metabolism were among the most significantly upregulated proteins in *CHD4* KD cells (Fig. 4A). This included MCAM, BCAM, ALCAM, COL1A1, COL1A2, CNN1, CKB, GPD1 and ALPL. Importantly, whereas RIPK3 and CNN1 were too low to be quantified by mass spectrometry in unperturbed cells (scramble shRNAs), their LFQ intensities increased to the upper 50^th^ percentile in CHD4-depleted cells (Fig. 4B). Thus, while CHD4 exerts genome-wide influence on gene expression, a group of genes including *RIPK3* exhibit an exceptional sensitivity to CHD4-mediated repression.

**Figure 4.**
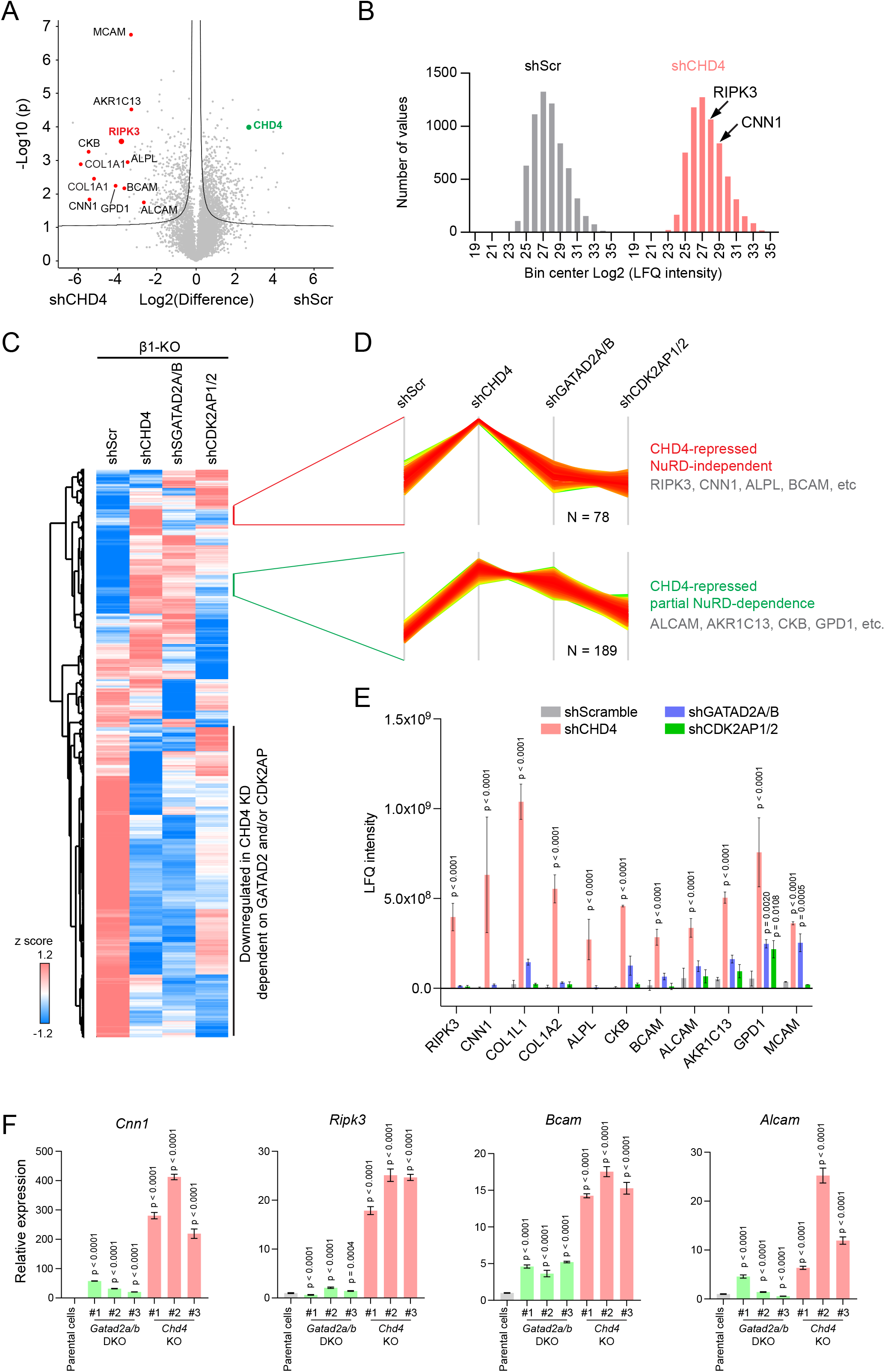
CHD4 shows high locus specificity independent of NuRD and ChAHP complexes. **(A)** Volcano plot of differential protein expression between β1-KO cells expressing control shScr and CHD4 shRNA. CHD4 and outliers that were most significantly upregulated in CHD4-depleted cells, including RIPK3, are highlighted (n = 3; FDR = 0.05, S0 = 0.1). **(B)** Histogram of logarithmically transformed LFQ intensities of the total proteome in β1-KO cells expressing control shScr and CHD4 shRNA (bin width = 2). The positions of RIPK3 and CNN1 are indicated by arrows. **(C)** Heatmap showing z-score of mean LFQ intensities of 2,833 differentially expressed proteins in control β1-KO, CHD4-depleted, GATAD2A/B-depleted and CDK2AP1/2-depleted cells. **(D)** A cluster of 78 proteins strongly upregulated in CHD4-depleted cells but not in GATAD2A/B-or CDK2AP1/2-depleted cells, including RIPK3, CNN1, ALPL, BCAM, etc., and another cluster of 189 proteins preferentially upregulated in CHD4-depleted cells, including ALCAM, AKR1C13, GPD1, CKB, etc., were identified. **(E)** Untransformed LFQ intensities of the most significantly upregulated proteins in CHD4-depleted cells were compared between cell lines depleted of GATAD2A/B or CDK2AP1/2. Data are depicted as mean ± s.d. (n = 3); p values were calculated with a two-way ANOVA with Tukey correction for multiple testing (for each protein, a comparison between the shScramble condition and the other knockdown conditions was conducted). **(F)** qPCR analysis of *Cnn1*, *Ripk3*, *Bcam* and *Alcam* transcript expression in parental β1 KO cells, three *Chd4* KO clones and three *Gatad2a/b* DKO clones using Gapdh as normalisation. Data are depicted as mean ± s.d. (n = 3); p values were calculated on log-transformed data by a one-way ANOVA with Tukey correction for multiple testing (for every cell line a comparison with the parental cells was conducted).

Whole proteome analysis of CHD4-depleted cells, GATAD2A/B-depleted cells and CDK2AP1/2-depleted cells revealed that 2,831 proteins among 6,498 quantified proteins were significantly changed in at least one of the cell lines (Fig. 4C, Supplementary Table 5). Notably, z-score clustering analysis revealed one group of 78 proteins strongly upregulated in CHD4-depleted cells but not in GATAD2A/B-or CDK2AP1/2-depleted cells, and another group of 189 proteins preferentially upregulated in CHD4-depleted cells (Fig. 4D). Surprisingly, the most significantly upregulated proteins in CHD4-depleted cells including RIPK3, CNN1, COL1A1, BCAM, ALPL, ALCAM, CKB, GPD1 belong to these two clusters (Fig. 4D & E). In contrast, none of these proteins were significantly upregulated in ADNP-depleted cells (Fig. S4A, Supplementary Table 6). To confirm that the changes in protein levels were due to transcriptional regulation, we performed qPCR analysis of a subset of CHD4-repressed genes in *Chd4* KO and *Gatad2a/b* DKO clones. As expected, the transcripts of *Cnn1*, *Ripk3*, *Bcam* and *Alcam* were dramatically upregulated in *Chd4* KO cells (Fig. 4F). *Cnn1* and *Bcam* were also slightly upregulated in *Gatad2a/b* DKO cells (Fig. 4F) likely due to reduced CHD4 protein levels in these cells (Fig. 3D). In contrast to the large changes in proteomics (Fig. 4E), *Col1a1* transcripts showed only 2-3-fold upregulation in the three *Chd4* KO clones (Fig. S4B). We took advantage of the collagen binding-deficient β1-KO cells and analyzed protein expression of collagen type 1 by immunostaining. This experiment revealed a subpopulation (around 20%) of CHD4 knockdown cells with dramatic intracellular accumulation of collagen type 1 protein (Fig. S4C). Hence, the increased collagen 1 protein levels in CHD4-deficient cells may be due to transcriptional upregulation that is only operational in a subset of cells. In summary, CHD4 can potently suppress gene expression independent of NuRD or ChAHP complexes with supreme locus specificity.

### Identification of CHD4-responsive CRE controlling RIPK3 expression

Previous ChIP-PCR studies of CHD4 have identified regions around the *Ripk3* locus that are enriched for binding of CHD4, HDAC1 and acetylated histone 3 *(25, 29)*. However, our data showed that neither NuRD nor HDAC1/2 activity is required for CHD4-mediated RIPK3 repression. Moreover, widespread genome binding of NuRD and ChAHP complexes does not permit precise determination of CHD4-responsive CREs controlling RIPK3 expression. To resolve this issue, we performed ATAC-seq experiments of the parental β1-KO cells, three individual *Chd4* KO clones and three individual *Gatad2a/b* DKO clones. We further included three biological replicates of β1-KO cells infected with a scramble shRNA virus to control random chromatin alteration due to genetic perturbation (ATAC-seq descriptive statistics in Supplementary Table 7). Heat map of z-score clustering (Fig. 5A) and PCA (Fig. S5A) analysis revealed that control cells, *Chd4* KO clones and *Gatad2a/b* DKO clones clustered into different groups each with robust yet distinct chromatin remodeling patterns. Among 74,164 retrieved ATAC-seq peaks, 17,284 were significantly increased and 7,969 were decreased in *Chd4* KO cells (Fig. 5B, Supplementary Table 8 & 9). In contrast, *Gatad2a/b* DKO cells showed less effect on chromatin accessibility with 5,012 ATAC-peaks increased and 2,143 decreased (Fig. 5C, Supplementary Table 10 & 11). Thus, both CHD4 and GATAD2A/B-containing NuRD complexes overall restrict chromatin accessibility. Approximately 4,000 increased and 1,600 decreased ATAC-seq peaks were found to be common for both *Chd4* KO and *Gatad2a/b* KO cells, which account for 73% significantly changed ATAC-seq peaks in *Gatad2a/b* DKO cells but only for 22% of that in *Chd4 KO* cells (Fig. 5D). Moreover, for ATAC-seq peaks commonly changed in *Chd4* KO and *Gatad2a/b* DKO cells and for peaks uniquely changed in one of the KO cells, the effect size was significantly higher in *Chd4* KO cells (Fig. 5E & F). Hence, although majority of GATAD2A/B-dependent chromatin openings relied on CHD4, CHD4 exerts much stronger and broader impact on chromatin accessibility independent of the NuRD complex.

**Figure 5.**
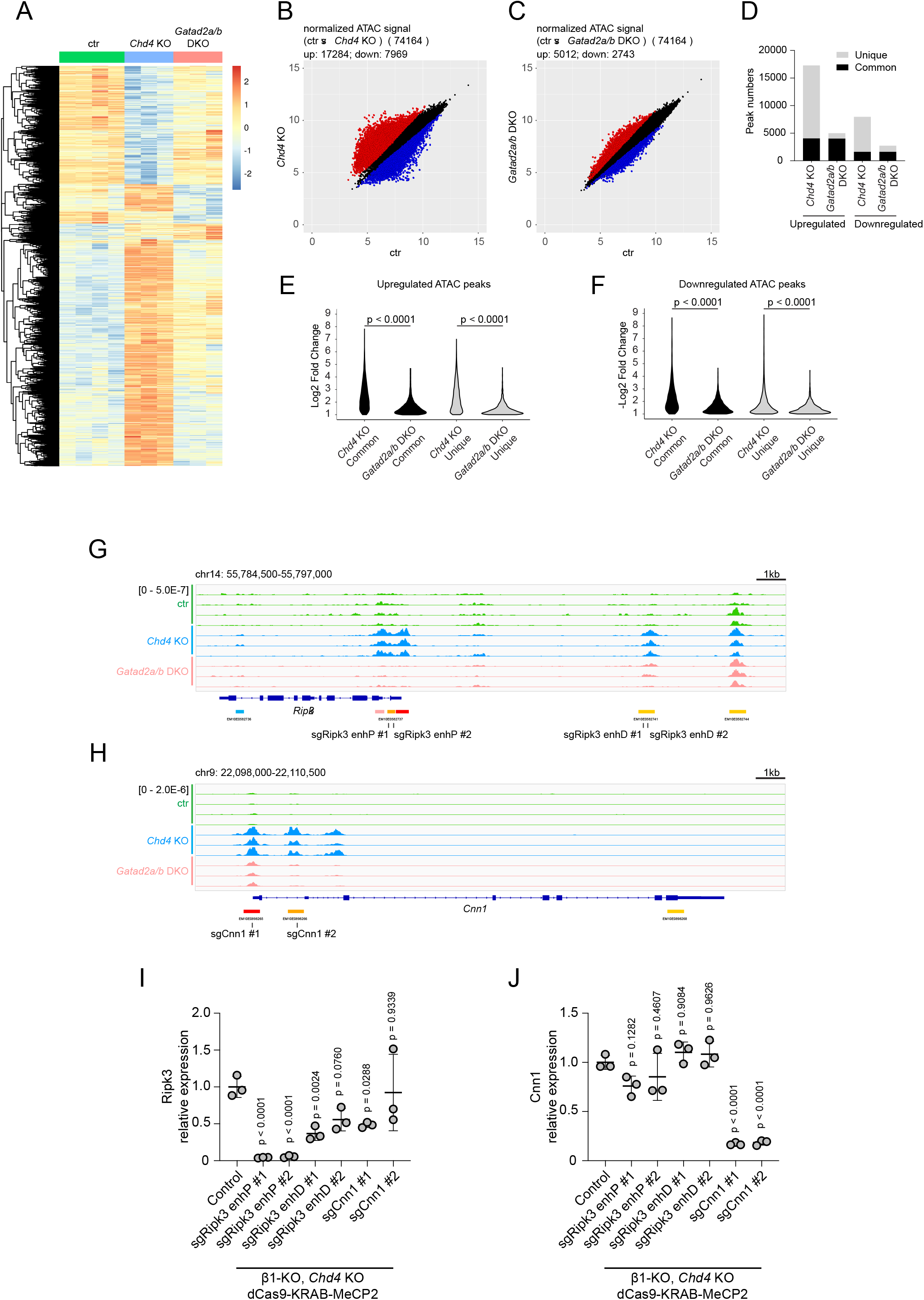
Identification of CHD4-responsive CRE controlling RIPK3 expression by ATAC-seq and CRISPR-silencing. (**A**) Heatmap clustering of differentially accessible regions (adjusted p-value <0.05 and log2 fold change > 1) between control, *Chd4* KO and *Gatad2a/b* DKO cells. **(B & C)** Scatter plot of differentially accessible regions between control cells and Chd4 KO cells **(B)** and between control cells and Gatad2a/b DKO cells **(C)**. Regions that gained or lost accessibility are highlighted in red and blue, respectively. **(D)** ATAC-seq peaks that were commonly altered in both *Chd4* KO cells and *Gatad2a/b* DKO cells, or uniquely altered in only one cell line, were identified and counted. **(E & F)** Violin plot showing the logarithmically transformed fold change of ATAC-seq signals of peaks commonly altered in both *Chd4* KO and *Gatad2a/b* DKO **(E)** or uniquely altered in one cell line **(F)**. p-values were calculated by Student t-test. **(G & H)** ATAC-seq signal traces near the *Ripk3* locus **(G)** and *Cnn1* locus **(H)**. ENCODE annotated cis-regulatory elements are highlighted. The positions and names of individual CRISPR sgRNAs are annotated below with black lines. **(I & J)** *Chd4* KO cells stably expressing dCas9-KRAB-MeCP2 were transduced with lentivirus expressing the indicated sgRNAs. mRNA expression of *Ripk3* **(I)** or *Cnn1* **(J)** in transduced cells and uninfected control cells was analysed by qPCR. Data are depicted as mean ± s.d. (n = 3) normalized to the mean value of the control condition; p values were calculated on log-transformed data by a one-way ANOVA with Dunnett correction for multiple testing (a comparison of the control cells with all other cell lines was conducted).

We next wished to functionally interrogate CHD4-responsive and GATAD2A/B-independent chromatin openings around the *Ripk3* locus using CRISPR-silencing in *Chd4* KO cells stably expressing dCas9-KRAB-MeCP2 *(45)*. Our ATAC-seq experiments had shown that there were no CHD4-responsive chromatin openings in regions previously identified by CHD4 ChIP-PCR around the *Ripk3* locus *(25, 29)*. However, we had observed a robust increase in chromatin accessibility around a distal enhancer and a proximal enhancer corresponding to EM10E0582741 and EM10E0582738 in ENCODE annotation, respectively (Fig. 5G). Two sgRNAs were stably expressed by lentiviral transduction to target the proximal (sgRipk3 enhP #1 and sgRipk3 enhP #2) and distal (sgRipk3 enhD #1 and sgRipk3 enhD #2) enhancers. As an additional control, to rule out the nonspecific effects of viral transduction and potential sgRNA off-targets, we also targeted the promoter and enhancer elements of the *Cnn1* locus, which had also gained accessibility in *Chd4* KO cells (sgCnn1 #1 and sgCnn1 #2, Fig. 5H). While CRISPR-silencing of the proximal enhancer of *Ripk3* locus resulted in 95% reduction in *Ripk3* mRNA levels and markedly reduced RIPK3 protein levels, targeting the distal enhancer of the *Ripk3* locus had only moderate influence (around 50% reduction) on *Ripk3* mRNA expression with marginal statistical significance and no obvious reduction of RIPK3 protein expression (Fig. 5I & Fig. S5C). Although none of the *Ripk3*-tageting sgRNA affected *Cnn1* transcript expression, both *Cnn1*-targeting sgRNA reduced *Cnn1* transcript by 80% with minimal influence on *Ripk3* expression (Fig. 5I & J). To test whether *Ripk3* expression is influenced by more distant CHD4-responsive elements via long-range genomic interactions, we also targeted other CHD4-responsive chromatin opening regions within 150kb range around *Ripk3* locus, most notably in nearby *Adcy4* and *Nfat3* locus (Fig. S5B). However, none of these sgRNAs altered RIPK3 protein levels (Fig. S5C). Taken together, our results indicated that the CHD4-responsive proximal enhancer of the *Ripk3* locus is the primary driver of high *Ripk3* gene expression.

## Discussion

Elevated RIPK3 expression is frequently observed in tissue fibrosis and may contribute to the sustained TNFα-driven inflammation in associated diseases. Studying RIPK3 expression in BDL-induced liver fibrosis, we could confirm a strong increase of RIPK3 expression in fibrosing tissue, yet we made the unexpected finding of RIPK3 being primarily induced in myofibroblasts. This contrasts previous literature, where RIPK3 expression has mainly been attributed to the myeloid compartment. Intrigued by this finding, we set out to dissect the mechanism of enhanced RIPK3 expression in a reductionist *in vitro* setting. Hypothesizing that cell adhesion to fibrotic ECM components would be involved in this regulation, we turned our attention to the β1 integrin receptor that is required for collagen and fibronectin binding. Interestingly, loss of β1 integrin in fibroblasts not only abolished ECM fibrillogenesis but also led to epigenetic repression of *Ripk3* locus by the chromatin-remodeling factor CHD4. Surprisingly this regulation displayed exquisite locus specificity and occurred independently of the conventional CHD4-associated NuRD or ChAHP complexes. Our data indicate that β1 integrin receptor intrinsically links fibrotic signaling to RIPK3-driven inflammation via a novel mode of action of CHD4.

Precise transcriptional regulation requires concerted, locus-specific actions of transcription factors and epigenetic modifiers. Class-II Chromodomain helicase DNA-binding (CHD) proteins including CHD3, CHD4 and CHD5 are associated with two distinct chromatin remodeling complexes, NuRD (Nucleosome Remodeling Deacetylase) and ChAHP. Emerging evidence suggest that CHD4 may also function outside the NuRD complex. For instance, CHD4 is only transiently associated with the rest of the pre-assembled NuRD subunits *in vitro* and processes nucleosome-remodeling activity by itself *(41)*. Mi-2β, the CHD4 homologue in Drosophila, suppresses germline gene expression in neurons independently of NuRD through a distinct complex containing Mep-1 *(46)*. However, there is no Mep-1 homologue in vertebrates. Here, we provide definitive evidence that CHD4 can regulate gene expression independently of the NuRD or ChAHP complexes with supreme locus specificity and potency in mammalian cells. Knockout or knockdown of CHD4, GATAD2A/B and ADNP, uncoupled chromatin remodeling, ATPase activity and locus specificity of CHD4 from NuRD or ChAHP complexes. Strikingly, the most significantly de-repressed genes in CHD4-depleted cells, which included *Ripk3*, *Cnn1*, *Bcam*, *Alcam*, etc., were not regulated by ChAHP or NuRD. Due to the interdependence of protein stability between subunits within NuRD or ChAHP complexes *(47)*, depletion of GATAD2A/B resulted in an approximately 50% reduction of CHD4 protein level, which explains why some NuRD-independent CHD4 target genes are mildly perturbed by GATAD2A/B depletion.

ATAC-seq analysis revealed much stronger and broader alterations of chromatin accessibility in *Chd4* KO in comparison with *Gatad2a/b* DKO. Whereas majority of chromatin openings in *Gatad2a/b* DKO cells are CHD4-dependent, 73% of CHD4-dependent chromatin openings do not depend on GATAD2A/B. Considering the 50% reduction of CHD4 in GATAD2A/B-depleted cells, the actual impact of NuRD-independent function of CHD4 may be even larger. NuRD-independent functions of CHD4 may be contributed by the ChAHP complex. However, ADNP depletion caused only 20% reduction of CHD4 (Fig. 3G), suggesting that only a small portion of CHD4 is engaged in the ChAHP complex in our cells. Furthermore, it has been reported that only around 30% CHD4-dependent ATAC-seq loci overlap with ADNP-dependent loci *(48)*. Therefore, CHD4 has profound impact on chromatin accessibility and gene expression independent of NuRD or ChAHP. Such a ‘stand-alone’ function may not be revealed by ChIP-seq analysis of CHD4 and other NuRD or ChAHP subunits since they may be passively recruited to CHD4-bound loci due to their high affinity interaction with CHD4. Our observations thus call for caution in using CHD4/NuRD or CHD4/ChAHP as equivalent to CHD4 in the future. CHD4 binds nucleosomes with no DNA sequence preference and its locus-targeting specificity relies on accessory transcription factors. Besides the high affinity interactions with NuRD and ChAHP complexes, CHD4 may bind epigenetic marks and transiently associate with a plethora of TFs to target to diverse genomic loci. Although class-II CHDs can all participate in NuRD or ChAHP complexes, they non-redundantly regulate distinct genes in neurodevelopment *(36)*. Their ‘stand-alone’ function may contribute to their diverse locus specificity and non-redundant functions *in vivo*. For instance, we observed that RIPK3 de-repression upon CHD4 depletion in mouse fibroblasts could be reversed by human CHD4 but not by human CHD5.

As one of the most prominent targets of the ‘stand-alone’ function of CHD4, RIPK3 plays an essential role in TNFα signaling and necroptosis. RIPK3 upregulation is frequently reported in liver fibrosis where it may not only promote necroptosis but also drive the expression of pro-inflammatory cytokines that further worsen disease progression. However, despite the beneficial effect of RIPK3 inhibition in various liver disease models, it remains controversial whether RIPK3-mediated necroptosis is relevant in liver cell death due to its low expression in hepatocytes *(26, 49)*. In bile duct ligation-induced liver fibrosis, we observed high RIPK3 expression in myofibroblasts, macrophages but not in hepatocytes. Since myofibroblasts are the main culprits in driving excessive ECM deposition and tissue stiffening in fibrosis, we erased the fibrotic signature in *in vitro* cultured fibroblasts by genetic ablation of β1 integrin. This not only abolished ECM fibrillogenesis but also led to a progressive loss of RIPK3 expression. Conversely, using a well-established integrin pan-KO fibroblast cell model, we confirmed that reconstitution of β1 integrin drove high RIPK3 expression. Importantly, diminished RIPK3 expression in β1 integrin KO cells could be reverted by CHD4 depletion, suggesting that CHD4-mediated repression of the RIPK3 locus is relieved in myofibroblasts downstream of profibrotic β1 integrin signaling. Previous ChIP-PCR data suggested that NuRD may be recruited to the *Ripk3* locus in regions around -1kbp and -6kbp upstream of the transcription start site. However, the widespread chromatin binding of NuRD and ChAHP does not permit precise definition of the CRE responsible for RIPK3 induction. We thus targeted multiple CHD4-responsive CREs within 150kbp range around *Ripk3* locus through CRISPR-silencing and defined proximal enhancer within *Ripk3* locus as the critical CRE driving *Ripk3* expression. This result may explain the hierarchical regulation of *Ripk3* by DNMT1 and CHD4 around its promoter.

Integrin receptors can elicit pleiotropic signaling pathways depending on receptor subtypes. For instance, the downstream signaling of α5β1 and αVβ3 integrins differ qualitatively, quantitatively, and kinetically in terms of the involvement of extracellular-signal regulated kinases (ERK), focal adhesion kinase (FAK), p21-activated kinases (PAK), Gα12/13, Rho family GTPase, and other factors. The unique combinatorial signaling pattern of β1 integrin might be relayed to the *Ripk3* locus via a yet unknown CHD4-containing complex. This β1 integrin-directed regulation could be fundamentally different from classical mechano-transduction pathways such as MKL1-SRF and YAP/TAZ that are synergistically activated by α5β1 and αVβ3 integrins. Due to the essential role of β1 integrin in the mesenchyme to maintain tissue integrity, genetic ablation of β1 integrin in fibroblasts *in vivo* will likely cause complex phenotypes and hence was not pursued in this study. However, our *in cellulo* experiments do point to an intrinsic connection between profibrotic β1 integrin signaling and RIPK3-dependent inflammation. Regulation of RIPK3 expression by β1 integrin signaling may be an integral part of the wound healing response to provide general immune protection in structural cells and to promote the release of growth factors for tissue regeneration *(50, 51)*. Given the key functions of RIPK3 in inflammation and tumor immunology, targeting the ‘stand-alone’ function of CHD4 may be a promising strategy to treat fibro-inflammation and to reactivate necroptosis in cancer cells. The set of CHD4-responsive genes defined in this study will allow genetic and chemical screening of novel CHD4 regulators in the future.

## Data deposition

Raw MS data is deposited in PRIDE database under accession number PXD041505. ATAC-seq data will be deposited in the GEO database (https://www.ncbi.nlm.nih.gov/geo/).

## Author contribution

Z.S. and V.H. are responsible for the conceptualization of the study, data curation and analysis, writing and editing of the manuscript. Z.S. performed most of the experiments. F.M.C. carried out the ATAC-seq experiments. F.M.C. and G.S. performed the ATAC-seq data analysis. H.H. and Z.A. provided mouse tissues samples of BDL model. A.Y. performed RNA-seq analysis.

## Acknowledgements

We thank Dr. Barbara Steigenberger (MPI Biochemistry) and MS core facility of Max Planck Institute of Biochemistry for MS proteomic analysis. We are grateful to Prof. Reinhard Fässler for generous support with critical cell models and research infrastructure in MPI Biochemistry. We thank Dr. Emanuel Rognoni and Dr. Michaela-Rosemarie Hermann for sample collection of RNA-seq analysis. We thank Dr. Mohammad Rahbari and Prof. Mathias Heikenwälder for insightful discussion and suggestions. This work is supported by the Deutsche Forschungsgemeinschaft (DFG, German Research Foundation) CRC 1403 (project number 414786233) to V.H. and TRR237 (project number 369799452) to V.H. and Z.A. Work in the G.S. lab is funded by the European Joint Programme on Rare Diseases (grant number EJPRD20-191).

## Materials and methods

### Animals and bile duct ligation model

8-9 week old mice C57BL/6J (B6) were purchased from Janvier (Le Genest-Saint-Isle, France. Liver fibrosis was induced in male, 8-9week old mice via bile duct ligation (BDL). Briefly, animals were treated with painkillers and anesthetized before the peritoneal cavity was opened along the linea alba. Two ligatures were placed around the common bile duct to obstruct it, before the incisions in the peritoneum and the skin were then closed, and the mice were allowed to recover. During the first 5 days after operation, all animals received additional injections of painkillers and liver injury was allowed to develop for 10 days before organs were harvested. Control mice were sham-operated (no ligation of the bile duct). All mice were kept according to the guidelines of the Federation of Laboratory Animal Science Association and experiments were authorized by permission of the LANUV in the state of Northrhine Westfalia (AZ. 84-02.04.2017.A222).

### Antibodies

See Supplementary Table 1.

### Plasmids DNA

pBABE-mRIPK3 was acquired from Addgene (pBabe-puro-mRipk3, Addgene plasmid #78830). hCHD5 cDNA was purchased from Addgene (pENTR3c GW CHD5, Addgene plasmid #68869). hCHD4 cDNA was a gift from Lucas Farnung. cDNA encoding HA-tagged human CHD4 and CHD5 were amplified by PCR and inserted into pB-EF-Bos vector *(52)*. To generate chimeric protein of N-terminus of hCHD5 fused to C-terminus of hCHD4, DNA encoding a.a.1-369 in hCHD4 was replaced by DNA encoding a.a.1-342 in hCHD5. Stable knockdown was achieved by lentivirus. shRNAs were targeting mouse CHD4 (shCHD4 #1: 5’-GCCCATCTTCTGAGTTGTAAA-3’; shCHD4 #2: 5’-TCGAGTGAGGACGACGATTTA-3’; shCHD4 #3: 5’-CGTAAACAGGTCAACTACAAT-3’), GATAD2A (5’-GTGCTGCCAATAATGAGTTTA-3’), GATAD2B (5’-ACAGGAAATTGAACAGCGATT-3’), CDK2AP1 (5’-GAAAGAGATCAGACCGACGTA-3’), CDK2AP2 (5’-TGGCAGAGACAGAACGCAATG-3’) and ADNP (5’-GCCTACAGATACCCTACTCAA-3’) were cloned into pLKO.1 vector. A scramble shRNA-expressing pLKO.1 (Addgene Plasmid #1864) was used as control (shScr). To generate knockout cells through CRISPR-Cas9, sgRNA targeting mouse *Chd4* (5’-TCTTACGGCTCCGACTACTG-3’), *Gatad2a* (5’-GGGCGCACGAACCTGAAGTG-3’), *Gatad2b* (5’-CGTCTAGCACTCATATCCAC-3’) were cloned into pSpCas9(BB)-2A-Puro (PX459) V2.0 *(53)* (Addgene plasmid #62988). SgRNAs for CRISPR silencing (sgRipk3 enhP #1: 5’-GAAACCCGACGTCTACGGCT-3’; sgRipk3 enhP #2: 5’-TGTCTCCGGCACACCCACAC-3’; sgRipk3 enhD #1: 5’-ACGACAATCTGACACTTTGG-3’; sgRipk3 enhD #2: 5’-AGGAGACCTAAGTTGCTGCC-3’; sgCnn1 #1: 5’-GCCTGTCCCATTGGCCACGG-3’; sgCnn1 #2: 5’-GGCCCGCGCTATATAAGGGC-3’; sgAdcy4: 5’-GCGGGTACAGAAGTAAACCG-3’; sgNfat3 #1: 5’-GCGTGACAGTGATCTACAGT-3’; sgNfat3 #2: 5’-CTCCCGGTTTCAATCACCTG-3’; sgNfat3 #3: 5’-GTTTGTAAACGTCTGACCAG-3’; sgNfat3 #4: 5’-GTCGAAGGGTGATGCGTGCG-3’; sgNfat3 #5: 5’-GAAGCCATTTGCAAGAACCT-3’; sgNfat3 #6: 5’-GATGTCCCATGGGATAAGGG-3’; sgNfat3 #7: 5’-CACTGGGGCTTTCAAATAGG-3’; sgNfat3 #8: 5’-CAGTGATAGCTGGGAATCAC-3’) were cloned into LentiSingle vector via ligation-independent cloning.

### Transient and stable gene expression in cell lines

SV40 large T-immortalized mouse kidney-derived fibroblasts have been previously described *(4, 43)*. The cell lines used here were not found in the database of commonly misidentified cell lines maintained by ICLAC and NCBI BioSample. All cell lines were tested negative for mycoplasma contamination. Cells were cultured in DMEM medium (Gibco) with 10% FBS (Gibco) and 1% penicillin–streptomycin (Gibco) under sterile conditions at 5% CO2 and 95% humidity. Transient transfections were carried out with Lipofectamine 2000 (Invitrogen) according to the manufacturer’s protocol. Briefly, 40 pmol ON-TARGETplus Mouse Chd4 SMARTPool siRNA (Dharmacon) was transfected with 4 µl Lipofectamine 2000 for each well of 6-well plate. 72hours after transfection, cells lysates were collected for analysis. To generate knockout cells through CRISPR-Cas9, 2 µg plasmid PX459 plasmid DNA was transfected with 4 µl Lipofectamine 2000 for each well of 6-well plate into target cells. 24 hours after transfection, cells were selected with 2 µg/ml puromycin for 24 hours. Surviving cells were single cell cloned and knockout was screened and validated by Western blot. To generate stable cell lines via lentivirus, VSV-G pseudo-typed lentiviral vectors were produced by transient transfection of HEK293T cells. Viral particles were concentrated from cell culture supernatant as previously described *(54)* and used for infection. To generate cell line stably expressing N-terminally HA-tagged human CHD4, CHD5 and chimeric protein containing N-terminus of hCHD5 fused to hCHD4 (HA-CHD4^CHD5-N^), piggy bac plasmids encoding these proteins and plasmid encoding hyperactive piggy bac transposase were co-transfected into target cells and selected with 2 µg/ml puromycin for 1week. For CRISPR silencing, *CHD4* KO cells was similarly transfected with piggy bac plasmid encoding dCas9-KRAB-MeCP2 *(45)* (Addgene plasmid #110821) and selected with 10 µg/ml blasticidin for 1week then infected with lentiviruses expressing different guide RNAs under U6 promoter.

### Lactate dehydrogenase (LDH) release assay

LDH release was determined with CyQUANT™ LDH Cytotoxicity Assay kit (Thermo Fisher Scientific). Whole cell lysates were generated by adding 10 x lysis buffer to cell culture at 37 °C for 30 min. Total cell lysates and supernatants were cleared by centrifugation at 450 x g for 10 min. 20 µL of supernatants were mixed with 20 µL reaction reagent mix and incubated for 10-30 min at RT in the dark and measured on Tecan Spark20M microplate reader for absorbance at 490nm and 680nm. LDH activity is calculated by subtracting absorbance at 490nm with absorbance at 680nm. Cell culture media was used as blank control. Percentage of LDH release was calculated by dividing LDH activity in cell culture supernatant with LDH activity in whole cell lysate.

### Quantitative RT-PCR

RNA was extract from indicated cells using TRIzol^TM^ reagent. Extracted RNAs were diluted into 100 ng/µl concentration. 500ng total RNA was used for cDNA synthesis using iScript™ cDNA Synthesis Kit (Bio-Rad). cDNA synthesized from 5ng total RNA was used as template for qPCR with SYBR Green mater mix (Bio-Rad) using a PCR setting of 95 °C, 3 min; 95 °C, 30 sec; 58 °C, 20 sec; 72 °C, 15 sec for 45 cycles. All qPCR primers were designed by Integrated DNA Technologies IDT with Tm value between 57 °C-59 °C and are listed in Supplementary Table 2.

### Mass spectrometry proteomics

For sample preparation, lysed cell pellets (in 300 µl PreOmics Lyse buffer) were incubation at 95 °C for 2 min and subsequently sonicated using a Bioruptor Plus sonication system (Diogenode) for 10×30sec at high intensity. Samples were incubated once more at 95 °C for 2 min and sonicated. Then, samples were diluted 1:1 with water and digested for 1.5 hours at 37 °C with 1 µg of LysC and overnight at 37 °C with 1 µg trypsin (Promega). The peptide mixture was acidified with trifluoroacetic acid (Merck) to a final concentration of 1%, followed by desalting of the peptides via SCX StageTips. Samples were vacuum dried and re-suspended in 6 µl of buffer A (0.1% formic acid).

For LC-MS/MS data acquisition, the peptides (3 µl) were loaded onto a 30 cm column (inner diameter: 75 microns; packed in-house with ReproSil-Pur C18-AQ 1.9-micron beads, Dr. Maisch GmbH) via the autosampler of the Thermo Easy-nLC 1200 (Thermo Fisher Scientific) at 60°C. Eluting peptides were directly sprayed onto the mass spectrometer Q Exactive HF (Thermo Fisher Scientific). Peptides were separated with a flow rate of 250nl/min by a gradient of buffer B (80% ACN, 0.1% formic acid) from 2% to 30% buffer B over 120 min followed an increase to 60%B over 10 min then to 95%B over the next 5 min and finally the percentage of buffer B was maintained at 95% buffer B for another 5 min. The mass spectrometer was operated in a data-dependent mode with MS1 scans from 300 to 1750 m/z (resolution of 60000 at m/z =200), and up to 15 of the top precursors were selected for fragmentation using higher energy collisional dissociation (HCD with a normalized collision energy of value of 28). The MS2 spectra were recorded at a resolution of 15000 (at m/z = 200). AGC target for MS1 and MS2 scans were set to 3×10^6^ and 1×10^5^ respectively within a maximum injection time of 100 and 25 ms for MS1 and MS2 scans respectively.

For data analysis, raw data were processed using the MaxQuant computational platform *(55)* (version 1.6.17.0). Briefly, the peak list was searched against the Uniprot database of Mus musculus with an allowed precursor mass deviation of 4.5 ppm and an allowed fragment mass deviation of 20 ppm. MaxQuant by default enables individual peptide mass tolerances, which was used in the search. Cysteine carbamidomethylation was set as static modification, and methionine oxidation and N-terminal acetylation as variable modifications. The match-between-run option was enabled, and proteins were quantified across samples using the label-free quantification algorithm in MaxQuant generating label-free quantification (LFQ) intensities.

Statistical analysis of MS measurements was performed in Perseus *(56)*. Briefly, raw data was filtered for proteins only identified by site, reverse peptide, and contaminants. Logarithmic transformation of LFQ intensities were filtered for valid values in 3 replicates in at least one cell types with imputation of missing values using default setting in Perseus (width = 0.3, down shift = 1.8). Volcano plot for comparison between two samples was calculated using standard student t-test (FDR = 0.05, S0 = 0.1). One-way ANOVA was used for statistical analysis of multiple comparison based on logarithmically transformed data (permutation-base FDR of 0.05). Significant targets were filtered, and z-score was calculated based on the mean value of their logarithmically transformed LFQ intensities. Original untransformed LFQ intensities were plotted in bar graphs.

### RNA-seq data analysis

After checking the quality of the samples (FastQC, v.0.10.1)*(57)*, cutadapt (v.1.4.1)*(58)* was used to remove the adapters and all reads which are afterwards were shorter than 80bp, followed by fastx_trimmer *(59)* (fastx toolkit, v. 0.0.14) to remove the first 10nt from the beginning of the reads (-f 11 -l 151 -Q33). When reads quality was assured, the files were mapped to the mouse genome (Genome build *NCBIM37*) downloaded from the iGenomes *(60)* using the tophat aligner (v. 2.0.11)*(61)* with default parameters for both single-and paired-end samples in the data set. The mapped files were then quantified on a gene level based on the ensembl annotations, using the featureCounts *(62)* (v. 1.4.2) tool from the SubRead package *(63)* (v. 2.0.1). Using the DESeq2*(64)* package (R 3.0.3*(65)*, DESeq2 version 1.2.10) the count data was normalized by the size factor to estimate the effective library size. This followed by the calculation of gene dispersion across all samples.

Perseus was used to perform statistical analysis of the RNA count data. Briefly, normalized DESeq2 count data was logarithmically transformed and filtered for valid values found in 3 replicates in at least one cell types with imputation of missing values using default setting in Perseus (width = 0.3, down shift = 1.8). Volcano plot for comparison between two samples was calculated using standard student t-test (FDR = 0.05, S0 = 1).

### ATAC-seq library preparation and data analysis

Omni-ATAC was performed as previously described *(66)* with minor modifications *(67)*. Briefly, 50000 cells were washed in 1xPBS, resuspended in 50μl of ATAC-seq resuspension buffer (RSB: 10 mM Tris-HCl, pH 7.4, 10 mM NaCl, and 3 mM MgCl2) containing 0.1 % NP40, 0.1 % Tween-20 and 0.01 % digitonin (Promega) and were incubated on ice for 3min. Following lysis, 1 ml of ATAC-seq RSB containing 0.1% Tween-20 was added, and nuclei were centrifuged at 500g (4 °C, 10 min). Pelleted nuclei were resuspended in 50 μl of transposition mix (25 μl 2 × TD buffer, 2.5 μl Tagment DNA enzyme (Illumina Tag DNA Enzyme and Buffer Kit, cat. 20034197), 16.5 μl PBS, 0.5 μl 1% digitonin, 0.5 μl 10% Tween-20, and 5.25 μl water) and incubated at 37 °C for 30 min in a thermomixer shaking at 1000 rpm. DNA was purified using Qiagen PCR clean-up MinElute kit (Qiagen). The transposed DNA was subsequently amplified in 50 µl reactions with custom primers as described *(68)*. After 4 cycles libraries were then monitored with qPCR: 5 µl PCR sample in a 15 µl reaction with the same primers. qPCR output was monitored for the ΔRN; 0.25 ΔRN cycle number was used to estimate the number of additional cycles of the PCR reaction needed for the remaining PCR samples. Amplified libraries were purified with the Qiagen PCR clean-up MinElute kit (Qiagen) and size selected for fragments shorter than 600 bp using the Agencourt AMPure XP beads (Beckman Coulter). Libraries were quality controlled by Qubit and Agilent DNA TapeStation analysis. Paired-end sequencing (60bp) was performed on an Illumina Next-Seq 2000 instrument.

The PEPATAC pipeline *(69)* (version 0.10.3) with default settings was used for primary analysis and quality controls. Briefly, after adapter trimming with Skewer *(70)* reads were first pre-aligned with bowtie2*(71)* to the mouse_chrM2x mitochondrial genome (parameters: -k 1 -D 20 -R 3 -N 1 -L 20 -i S,1,0.50) to remove mitochondrial reads and then mapped to the mm10 mouse genome (parameters: -- very-sensitive -X 2000). Following alignment, reads with mapping quality scores below 10 and any residual mitochondrial reads were removed and read deduplication was carried out using Samblaster *(72)*. Alignments were used to generate a signal track for visualization with IGV genome browser *(73)*. MACS2*(74)* was used for peak-calling (parameters: --shift -75 --extsize 150 --nomodel --call-summits -- nolambda --keep-dup all -p 0.01). Called peaks were filtered against the ENCODE blacklist *(75)* and were standardized by extending up and down 250 bases from the summit of each peak to establish peaks 500 bases in width.

To establish consensus peaks, overlapping (1 bp) peaks between every sample in the project were identified and the consensus peak’s coordinates based on the overlapping peak with the highest score were defined. Peaks present in at least 2 samples with a minimum score per million ≥5 were retained. A peak count table was then provided where every sample peak set is overlapped against the consensus peak set. Individual peak counts for an overlapping peak were weighted by multiplying by the percent overlap of the sample peak with the consensus peak. The consensus peak counts table was loaded into R *(65)* (version 4.1.0) and DESeq2*(76)* (version 1.32.0) was used for normalization and identification of significant differential accessible regions (adjusted p-value <0.05 and log2 fold change > 1). ggplot2*(77)* (version 3.3.5) and pheatmap *(78)* (version 1.0.12) were used data visualization. Samples mapping statistics and quality controls summary are provided in Supplementary Table 7.

### Immunofluorescence microscopy

For immunofluorescence of fixed cells, cells cultured on FN-coated (10μg/ml) coverslips were fixed with 4% PFA at room temperature for 20min before staining. For immunostaining of extracellular collagen type 3, cells were washed twice with PBS, blocked with 5% BSA in PBS for 1 hour and directly incubated with the primary antibody diluted in PBS at 4 °C overnight. For immunostaining of intracellular proteins, fixed cells were permeabilized with 0.1% Triton X-100 in PBS for 20min at room temperature. Fixed cells were blocked with 5% BSA in PBS for 1 hour at room temperature followed by incubation with the primary antibodies in 3% BSA in PBS overnight at 4 °C and then with fluorescently conjugated secondary antibodies for 1 h at room temperature. DNA was briefly stained with DAPI for 5min at room temperature. Images were collected at room temperature on a Zeiss (Jena) LSM780 confocal laser scanning microscope equipped with a Zeiss Plan-APO ×63–numerical aperture 1.46 oil immersion objective. For immunostaining of tissue samples, 7 µm thick cryosections were permeabilized with 0.05% Triton X-100 fin PBS for 30min at room temperature.

### Live cell imaging

3×10^5^ indicated cells were seeded on FN-coated (10μg/ml) plates overnight. Cells were treated with vehicle control, 40 µM zVAD, 50 ng/ml TNFα and 40 µM zVAD in combination as indicated in the presence of 5 µM Cytox^TM^Green (S7020, Invitrogen^TM^) and transferred to EVOS FL Auto live cell imaging system (Thermo Fisher Scientific) with environment control at 37 °C and 5% CO_2_. Temperature and focus were stabilized for 30min before brightfield and epifluorescence images were taken with 20X objective every 5min for 24hours. Fluorescent signals in time series images were measured by ImageJ.

### Immunoprecipitation

For immunoprecipitation of HA-tagged CHD proteins, indicated cells were lysed in IP buffer (50mM TrisHCl, pH 7.4, 150mM, 1% Triton X-100) with mild sonication. Supernatants of cell lysate were immunoprecipitated with and wash three time with IP buffer. Protein bound to the beads were eluted with Laemmli buffer at 95°C for 10min. Elutes were separated in SDS–PAGE for Western blotting.

### Statistics and reproducibility

Statistical analysis was carried out in GraphPad Prism software (version 9.00, GraphPad Software). Statistical calculations to predetermine required sample size were not carried out. Data sets were analysed using either Student’s t-tests, one-way ANOVA, or two-way ANOVA with additional corrections for multiple testing as indicated. In some graphs only the p values of relevant comparisons are shown. Results are depicted as mean ± s.d. or mean ± s.e.m. as indicated in figure legends. All experiments for quantitative analysis were reproduced at least three times. Source data and statistic test results for RNA-seq, proteomics and ATAC-seq are provided as Supplementary Tables. All other data that support the conclusions are available from the authors upon request.

## Supplementary figure legends

**Figure S1. RIPK3 expression in BDL-induced liver fibrosis**.

**(A)** 7 µm sections of cryopreserved liver tissue from sham-operated (Sham) or bile duct-ligated (BDL) mice were immunostained for smooth muscle actin (SMA, red), CD11b (blue), RIPK3 (green) and DAPI (grey channel in merged image). Images of healthy liver sections and fibrotic areas of the BDL model were obtained by confocal microscopy (scale bar 100 µm). **(B)** Zoomed image of fibrotic area of BDL liver section from the box indicated in **(A)** (scale bar 100 µm). **(C)** 7 µm sections of cryopreserved liver tissue from sham-operated (Sham) or bile duct-ligated (BDL) mice were immunostained for Vimentin (green), RIPK3 (red) and DAPI (blue). Images were taken by confocal microscopy (scale bar 100 µm). **(D)** Zoomed image of fibrotic area of BDL liver section from the box indicated in **(C)**. Fibroblasts positive for Vimentin and RIPK3 are indicated by the arrow and binucleated hepatocytes with low RIPK3 signal are indicated by the arrowhead (scale bar 40 µm). **(E & F)** The BDL liver section images were further segmented into fibrotic lesions and surrounding non-fibrotic areas in the BDL liver section. Fluorescence intensities of vimentin and RIPK3 were measured in the indicated areas. Data are depicted as mean ± s.d. (n = 7 from 3 pairs of operated mice); p values were calculated by one-way ANOVA with Tukey correction for multiple testing (a comparison of the sham condition with the different regions was conducted). **(G)** pKO-αV, pKO-αV/β1 and pKO-β1 cells were seeded on FN-coated (10 μg/ml) 6-well plates for 24 hours at confluence and treated with vehicle control or zVAD and 50 ng/ml TNFα in combination for a further 8 hours. *Cxcl10* and *Cxcl1* induction by necroptosis stimulus was analysed by qPCR and is depicted as mean ± s.d. (n = 3).

**Figure S2. CHD4 is the major gatekeeper of *Ripk3* expression in multiple cell lines**.

**(A)** pKO-αV, pKO-αV/β1 and pKO-β1 cells were seeded on FN-coated (10 μg/ml) 6-well plates and treated with 0.5 µM 5-Aza-2’-desoxycytidin (5’AZA) for 72 hours. *Ripk3* mRNA expression was analysed by qPCR and is depicted as mean ± s.d. (n = 3); p values were calculated on log-transformed data by a two-way ANOVA with a Tukey correction for multiple testing. **(B)** β1-WT and β1-KO cells were seeded on FN-coated (10 μg/ml) 6-well plates and treated with 0.5 µM 5-Aza-2’-desoxycytidin (5’AZA) for 72 hours. *Ripk3* mRNA expression was analysed by qPCR and is depicted as mean ± s.d. (n = 3); p values were calculated on log-transformed data by a two-way ANOVA with a Tukey correction for multiple testing. **(C)** β1-WT and β1-KO cells were seeded on FN-coated coverslips (10 μg/ml) for 24 hours and immunostained for CHD4 (green) and DAPI (blue). Scale bar, 20 μm. **(D)** The indicated cell lines are transiently transfected with non-targeting siScr or siRNA against CHD4 (siCHD4) for 72 hours. Protein expression of CHD4 and RIPK3 was analysed by immunoblot using VCL as a loading control. **(E)** RIPK3 protein levels in **(D)** were quantified by densitometric analysis and were normalized to β1-KO fibroblasts (siScr) are depicted as mean ± s.d (n = 3; only 2 data points for the *Chd4* KO fibroblasts); p values were calculated on log-transformed data using a paired t-test with Holm-Šídák’s correction for multiple testing.

**Figure S3. Perturbation of NuRD and ChAHP subunits by knockdown of CHD4, GATAD2A/B and CDK2AP1/2**.

LFQ intensity of individual NuRD and ChAHP subunits in β1-KO cells stably expressing control scramble shRNA, shRNA targeting GATAD2A/B or CDK2AP1/2. Due to the small size and high sequence homology of CDK2AP1 and CDK2AP2, these two paralogous proteins could not be distinguished by MS analysis. Data are depicted as mean ± s.d (n = 3); p values were calculated by one-way ANOVA with a Dunnett correction for multiple testing for each protein or paralogous group (a comparison of the different knockdown conditions with the respective shScr control was conducted).*, p < 0.05; **, p < 0.01; ***, p < 0.001; ****, p < 0.0001.

**Figure S4. CHD4 can regulate target gene expression independent of NuRD or ChAHP complexes**.

**(A)** LFQ intensity of selected proteins in β1-KO cells stably expressing control scramble shRNA or shRNA against ADNP. Data are depicted as mean ± s.d. (n = 3); p values were calculated by a two-way ANOVA with Šídák’s correction for multiple testing. **(B)** qPCR analysis of *Col1a1* mRNA expression in parental β1-KO cells, three *Chd4* KO clones and three *Gatad2a/b* DKO clones using Gapdh as normalisation. Data are depicted as mean ± s.d. (n = 3); p values were calculated on log-transformed data by one-way ANOVA with Tukey correction for multiple testing (a comparison of the parental cells with the other cell lines was conducted). **(C)** β1-KO cells stably expressing control scramble shRNA or shCHD4 were seeded on FN-coated coverslips (10 μg/ml) for 24 hours and immunostained for F-actin (phalloidin, red), collagen type 1 (green) and DAPI (blue). Scale bar, 20 μm.

**Figure S5. ATAC-seq and CRISPR-silencing analysis of CHD4-responsive cis-regulatory elements around *Ripk3* locus**.

**(A)** PCA analysis of ATAC-seq data from control cells (ctr, orange), *Chd4* KO cells (green) and *Gatad2a/b* DKO cells (grey) showed that three cell types grouped into distinct clusters. **(B)** ATAC-seq signal traces within a 67kb region around the Ripk3 locus. ENCODE annotated cis-regulatory elements are highlighted in colour. Positions and names of individual CRISPR sgRNAs are annotated below with black lines. **(C)** Chd4 KO cells stably expressing dCas9-KRAB-MeCP2 were transduced with lentivirus expressing the indicated sgRNAs. RIPK3 protein expression in transduced cells and uninfected control cells was analysed by immunoblot. Data are representative of 2 independent experiments.

## Supplementary data

Table S1. Antibodies used in the study.

Table S2. qPCR primers used in the study.

Table S3. DEseq2 count of RNA-seq data.

Table S4. Original LFQ intensities in MS measurement of shScr, shCHD4, shGATAD2A/B and shCDK2AP1/2 cells.

Table S5. One-way ANOVA test of Log transformed LFQ intensities shScr, shCHD4, shGATAD2A/B and shCDK2AP1/2 cells.

Table S6. Original LFQ intensities in MS measurement of shScr and shADNP cells.

Table S7. ATAC-seq sample statistics.

Table S8. ATAC-seq track data upregulated peaks in *Chd4* KO.

Table S9. ATAC-seq track data downregulated peaks in *Chd4* KO.

Table S10. ATAC-seq track data upregulated peaks in Gatad2a/b DKO.

Table S11. ATAC-seq track data downregulated peaks in Gatad2a/b DKO.

## References

1. N. C. Henderson, F. Rieder, T. A. Wynn, Fibrosis: from mechanisms to medicines. Nature 587, 555–566 (2020).

2. Z. Sun, M. Costell, R. Fassler, Integrin activation by talin, kindlin and mechanical forces. Nat Cell Biol 21, 25–31 (2019).

3. Z. Sun, S. S. Guo, R. Fassler, Integrin-mediated mechanotransduction. *J Cell Biol* **215**, 445–456 (2016).

4. H. B. Schiller et al., beta1- and alphav-class integrins cooperate to regulate myosin II during rigidity sensing of fibronectin-based microenvironments. Nat Cell Biol 15, 625–636 (2013).

5. M. R. Hermann et al., Integrins synergise to induce expression of the MRTF-A-SRF target gene ISG15 for promoting cancer cell invasion. J Cell Sci 129, 1391–1403 (2016).

6. X. He, et al., Myofibroblast YAP/TAZ activation is a key step in organ fibrogenesis. JCI Insight 7, (2022).

7. K. Martin et al., PAK proteins and YAP-1 signalling downstream of integrin beta-1 in myofibroblasts promote liver fibrosis. Nat Commun 7, 12502 (2016).

8. E. H. Danen, P. Sonneveld, C. Brakebusch, R. Fassler, A. Sonnenberg, The fibronectin-binding integrins alpha5beta1 and alphavbeta3 differentially modulate RhoA-GTP loading, organization of cell matrix adhesions, and fibronectin fibrillogenesis. J Cell Biol 159, 1071–1086 (2002).

9. S. He et al., Receptor interacting protein kinase-3 determines cellular necrotic response to TNF-alpha. Cell 137, 1100–1111 (2009).

10. Y. S. Cho et al., Phosphorylation-driven assembly of the RIP1-RIP3 complex regulates programmed necrosis and virus-induced inflammation. Cell 137, 1112–1123 (2009).

11. D. W. Zhang et al., RIP3, an energy metabolism regulator that switches TNF-induced cell death from apoptosis to necrosis. Science 325, 332–336 (2009).

12. S. He, X. Wang, RIP kinases as modulators of inflammation and immunity. Nat Immunol 19, 912–922 (2018).

13. L. Sun et al., Mixed lineage kinase domain-like protein mediates necrosis signaling downstream of RIP3 kinase. Cell 148, 213–227 (2012).

14. J. Zhao et al., Mixed lineage kinase domain-like is a key receptor interacting protein 3 downstream component of TNF-induced necrosis. Proc Natl Acad Sci U S A 109, 5322–5327 (2012).

15. F. Pinci et al., C-tag TNF: a reporter system to study TNF shedding. J Biol Chem 295, 18065–18075 (2020).

16. F. Pinci et al., Tumor necrosis factor is a necroptosis-associated alarmin. Front Immunol 13, 1074440 (2022).

17. Z. Cai et al., Activation of cell-surface proteases promotes necroptosis, inflammation and cell migration. Cell Res 26, 886–900 (2016).

18. K. Zhu et al., Necroptosis promotes cell-autonomous activation of proinflammatory cytokine gene expression. Cell Death Dis 9, 500 (2018).

19. L. S. Verjee et al., Unraveling the signaling pathways promoting fibrosis in Dupuytren’s disease reveals TNF as a therapeutic target. Proc Natl Acad Sci U S A 110, E928–937 (2013).

20. T. A. Wynn, T. R. Ramalingam, Mechanisms of fibrosis: therapeutic translation for fibrotic disease. Nat Med 18, 1028–1040 (2012).

21. J. E. Morgan et al., Necroptosis mediates myofibre death in dystrophin-deficient mice. Nat Commun 9, 3655 (2018).

22. M. B. Afonso et al., RIPK3 acts as a lipid metabolism regulator contributing to inflammation and carcinogenesis in non-alcoholic fatty liver disease. Gut, (2020).

23. A. Takezaki et al., A homozygous SFTPA1 mutation drives necroptosis of type II alveolar epithelial cells in patients with idiopathic pulmonary fibrosis. J Exp Med 216, 2724–2735 (2019).

24. M. Imamura, et al., RIPK3 promotes kidney fibrosis via AKT-dependent ATP citrate lyase. JCI Insight 3, (2018).

25. K. Sreenivasan et al., Attenuated Epigenetic Suppression of Muscle Stem Cell Necroptosis Is Required for Efficient Regeneration of Dystrophic Muscles. Cell Rep 31, 107652 (2020).

26. S. P. Preston et al., Epigenetic Silencing of RIPK3 in Hepatocytes Prevents MLKL-mediated Necroptosis From Contributing to Liver Pathologies. Gastroenterology 163, 1643–1657 e1614 (2022).

27. J. Gautheron et al., A positive feedback loop between RIP3 and JNK controls non-alcoholic steatohepatitis. EMBO Mol Med 6, 1062–1074 (2014).

28. G. B. Koo et al., Methylation-dependent loss of RIP3 expression in cancer represses programmed necrosis in response to chemotherapeutics. Cell Res 25, 707–725 (2015).

29. S. Colijn et al., The NuRD chromatin-remodeling complex enzyme CHD4 prevents hypoxia-induced endothelial Ripk3 transcription and murine embryonic vascular rupture. Cell Death Differ 27, 618–631 (2020).

30. J. K. K. Low et al., The Nucleosome Remodeling and Deacetylase Complex Has an Asymmetric, Dynamic, and Modular Architecture. Cell Rep 33, 108450 (2020).

31. V. Ostapcuk et al., Activity-dependent neuroprotective protein recruits HP1 and CHD4 to control lineage-specifying genes. Nature 557, 739–743 (2018).

32. M. Sharifi Tabar et al., Unique protein interaction networks define the chromatin remodelling module of the NuRD complex. FEBS J 289, 199–214 (2022).

33. M. Sharifi Tabar, J. P. Mackay, J. K. K. Low, The stoichiometry and interactome of the Nucleosome Remodeling and Deacetylase (NuRD) complex are conserved across multiple cell lines. FEBS J 286, 2043–2061 (2019).

34. J. G. Marques et al., NuRD subunit CHD4 regulates super-enhancer accessibility in rhabdomyosarcoma and represents a general tumor dependency. Elife 9, (2020).

35. N. Mor et al., Neutralizing Gatad2a-Chd4-Mbd3/NuRD Complex Facilitates Deterministic Induction of Naive Pluripotency. Cell Stem Cell 23, 412–425 e410 (2018).

36. J. Nitarska et al., A Functional Switch of NuRD Chromatin Remodeling Complex Subunits Regulates Mouse Cortical Development. Cell Rep 17, 1683–1698 (2016).

37. C. G. Spruijt et al., ZMYND8 Co-localizes with NuRD on Target Genes and Regulates Poly(ADP-Ribose)-Dependent Recruitment of GATAD2A/NuRD to Sites of DNA Damage. Cell Rep 17, 783–798 (2016).

38. A. Kumar et al., KSHV episome tethering sites on host chromosomes and regulation of latency-lytic switch by CHD4. Cell Rep 39, 110788 (2022).

39. F. Sher et al., Rational targeting of a NuRD subcomplex guided by comprehensive in situ mutagenesis. Nat Genet 51, 1149–1159 (2019).

40. J. K. Low et al., CHD4 Is a Peripheral Component of the Nucleosome Remodeling and Deacetylase Complex. J Biol Chem 291, 15853–15866 (2016).

41. W. Zhang et al., The Nucleosome Remodeling and Deacetylase Complex NuRD Is Built from Preformed Catalytically Active Sub-modules. J Mol Biol 428, 2931–2942 (2016).

42. L. Farnung, M. Ochmann, P. Cramer, Nucleosome-CHD4 chromatin remodeler structure maps human disease mutations. Elife 9, (2020).

43. R. T. Bottcher et al., Sorting nexin 17 prevents lysosomal degradation of beta1 integrins by binding to the beta1-integrin tail. Nat Cell Biol 14, 584–592 (2012).

44. Y. Zhong et al., The role of auxiliary domains in modulating CHD4 activity suggests mechanistic commonality between enzyme families. Nat Commun 13, 7524 (2022).

45. N. C. Yeo et al., An enhanced CRISPR repressor for targeted mammalian gene regulation. Nat Methods 15, 611–616 (2018).

46. G. N. Aughey, E. Forsberg, K. Grimes, S. Zhang, T. D. Southall, NuRD-independent Mi-2 activity represses ectopic gene expression during neuronal maturation. EMBO Rep, e55362 (2023).

47. T. Burgold et al., The Nucleosome Remodelling and Deacetylation complex suppresses transcriptional noise during lineage commitment. EMBO J 38, (2019).

48. X. Sun, W. Yu, L. Li, Y. Sun, ADNP Controls Gene Expression Through Local Chromatin Architecture by Association With BRG1 and CHD4. Front Cell Dev Biol 8, 553 (2020).

49. V. Kondylis, M. Pasparakis, RIP Kinases in Liver Cell Death, Inflammation and Cancer. Trends Mol Med 25, 47–63 (2019).

50. S. Zhou et al., Myofiber necroptosis promotes muscle stem cell proliferation via releasing Tenascin-C during regeneration. Cell Res 30, 1063–1077 (2020).

51. T. Krausgruber et al., Structural cells are key regulators of organ-specific immune responses. Nature 583, 296–302 (2020).

52. K. Yusa, L. Zhou, M. A. Li, A. Bradley, N. L. Craig, A hyperactive piggyBac transposase for mammalian applications. Proc Natl Acad Sci U S A 108, 1531–1536 (2011).

53. F. A. Ran et al., Genome engineering using the CRISPR-Cas9 system. Nat Protoc 8, 2281–2308 (2013).

54. A. Pfeifer, T. Kessler, S. Silletti, D. A. Cheresh, I. M. Verma, Suppression of angiogenesis by lentiviral delivery of PEX, a noncatalytic fragment of matrix metalloproteinase 2. Proc Natl Acad Sci U S A 97, 12227–12232 (2000).

55. J. Cox, M. Mann, MaxQuant enables high peptide identification rates, individualized p.p.b.-range mass accuracies and proteome-wide protein quantification. Nat Biotechnol 26, 1367–1372 (2008).

56. S. Tyanova et al., The Perseus computational platform for comprehensive analysis of (prote)omics data. Nat Methods 13, 731–740 (2016).

57. S. Andrews.

58. M. Martin, Cutadapt removes adapter sequences from high-throughput sequencing reads. EMBnet.journal 17, 3 (2011).

59. G. J. Hannon. (2010). Illumina, iGenomes (https://support.illumina.com/sequencing/sequencing_software/igenome.html).

60. D. Kim et al., TopHat2: accurate alignment of transcriptomes in the presence of insertions, deletions and gene fusions. Genome Biol 14, R36 (2013).

61. Y. Liao, G. K. Smyth, W. Shi, featureCounts: an efficient general purpose program for assigning sequence reads to genomic features. Bioinformatics 30, 923–930 (2014).

62. Y. Liao, G. K. Smyth, W. Shi, The Subread aligner: fast, accurate and scalable read mapping by seed-and-vote. Nucleic Acids Res 41, e108 (2013).

63. S. Anders, W. Huber, Differential expression analysis for sequence count data. Genome Biol 11, R106 (2010).

64. R: A language and environment for statistical computing. R Foundation for Statistical Computing, Vienna, Austria., (R Development Core Team, 2018).

65. M. R. Corces et al., An improved ATAC-seq protocol reduces background and enables interrogation of frozen tissues. Nat Methods 14, 959–962 (2017).

66. F. M. Cernilogar et al., Pre-marked chromatin and transcription factor co-binding shape the pioneering activity of Foxa2. Nucleic Acids Res 47, 9069–9086 (2019).

67. A. Mezger et al., High-throughput chromatin accessibility profiling at single-cell resolution. Nat Commun 9, 3647 (2018).

68. J. P. Smith, et al., PEPATAC: an optimized pipeline for ATAC-seq data analysis with serial alignments. *NAR Genom Bioinform* 3, lqab101 (2021).

69. H. Jiang, R. Lei, S. W. Ding, S. Zhu, Skewer: a fast and accurate adapter trimmer for next-generation sequencing paired-end reads. BMC Bioinformatics 15, 182 (2014).

70. B. Langmead, S. L. Salzberg, Fast gapped-read alignment with Bowtie 2. Nat Methods 9, 357–359 (2012).

71. G. G. Faust, I. M. Hall, SAMBLASTER: fast duplicate marking and structural variant read extraction. Bioinformatics 30, 2503–2505 (2014).

72. J. T. Robinson et al., Integrative genomics viewer. Nat Biotechnol 29, 24–26 (2011).

73. Y. Zhang et al., Model-based analysis of ChIP-Seq (MACS). Genome Biol 9, R137 (2008).

74. H. M. Amemiya, A. Kundaje, A. P. Boyle, The ENCODE Blacklist: Identification of Problematic Regions of the Genome. Sci Rep 9, 9354 (2019).

75. M. I. Love, W. Huber, S. Anders, Moderated estimation of fold change and dispersion for RNA-seq data with DESeq2. Genome Biol 15, 550 (2014).

76. H. Wickham, in Use R!,. (Springer International Publishing : Imprint: Springer,, Cham, 2016), pp. 1 online resource (XVI, 260 pages 232 illustrations, 140 illustrations in color.

77. R. Kolde. (2019).

