## Supplementary figures and images for "β1 integrin signaling governs necroptosis via the chromatin remodeling factor CHD4"

### Supplemental Figures 1-5

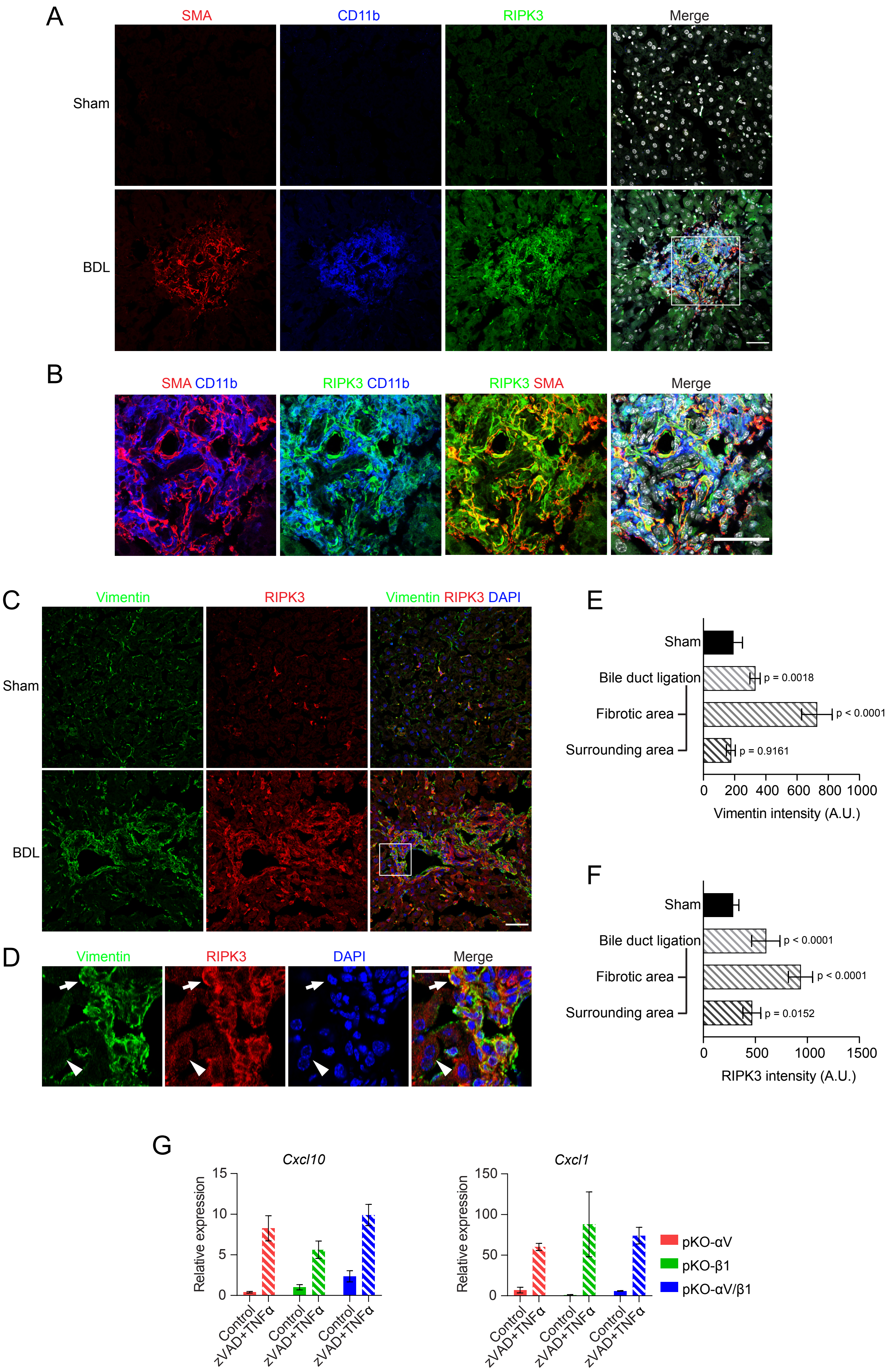

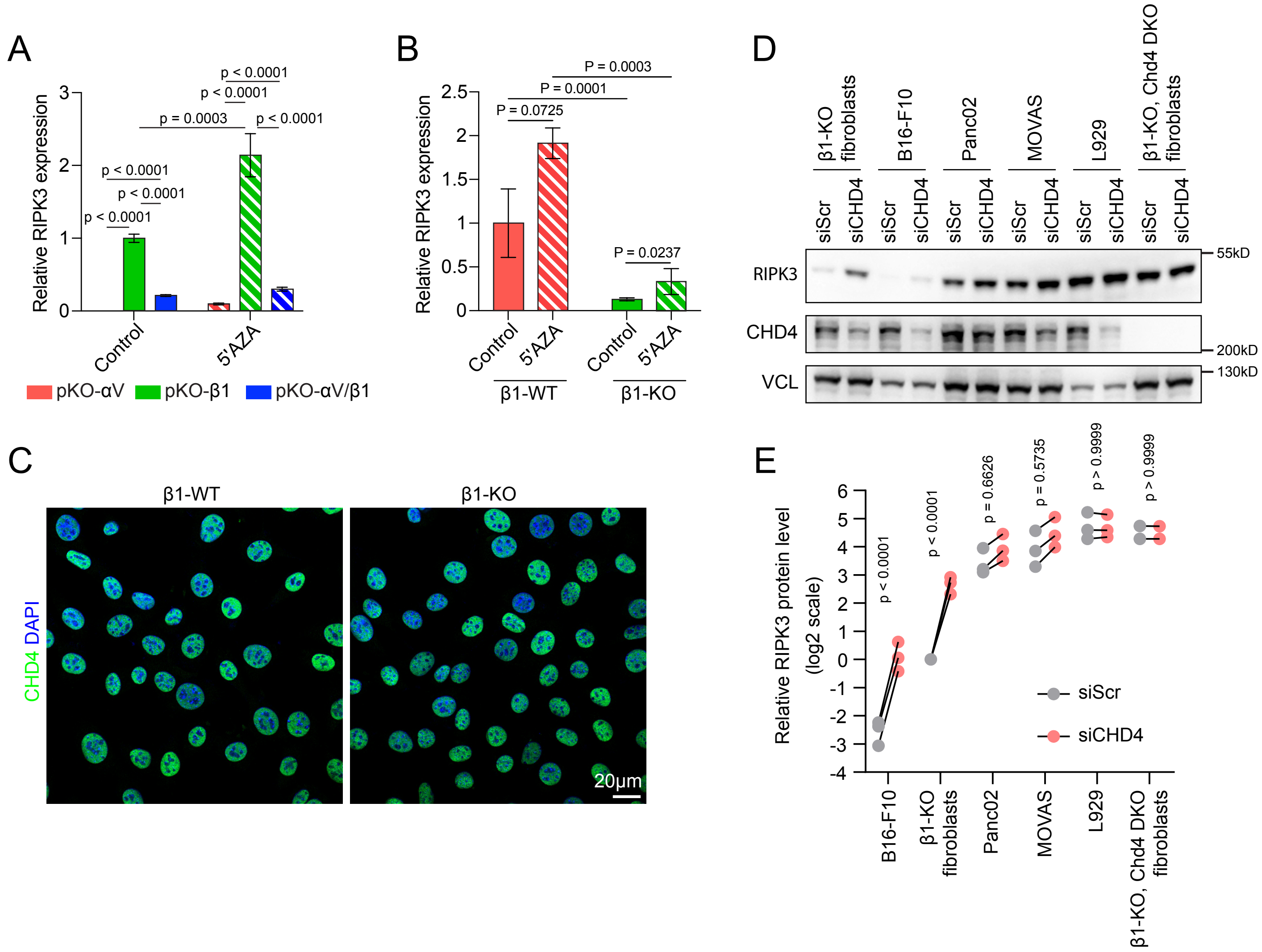

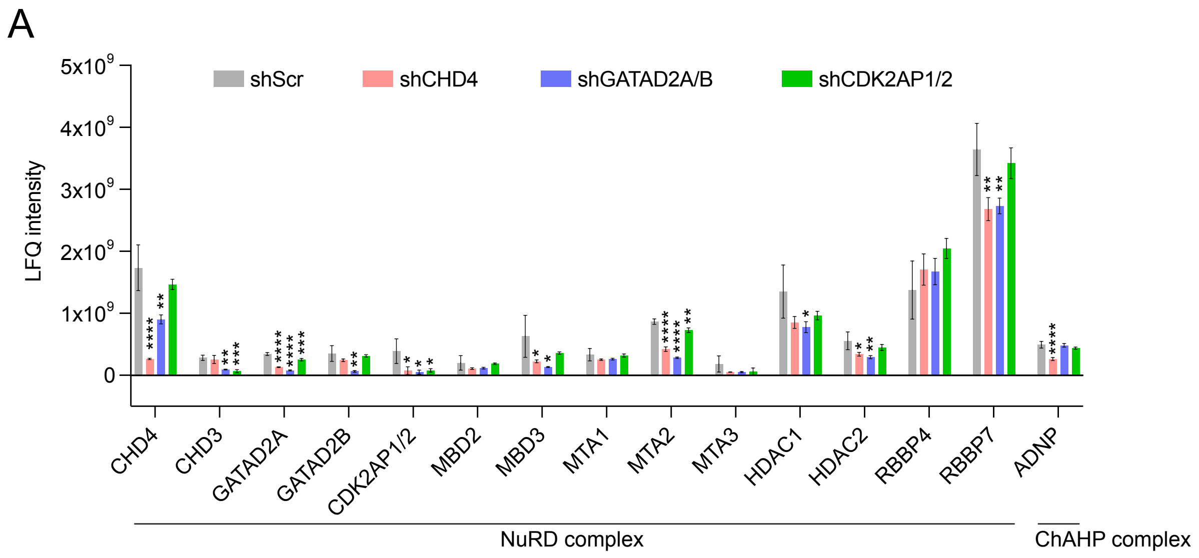

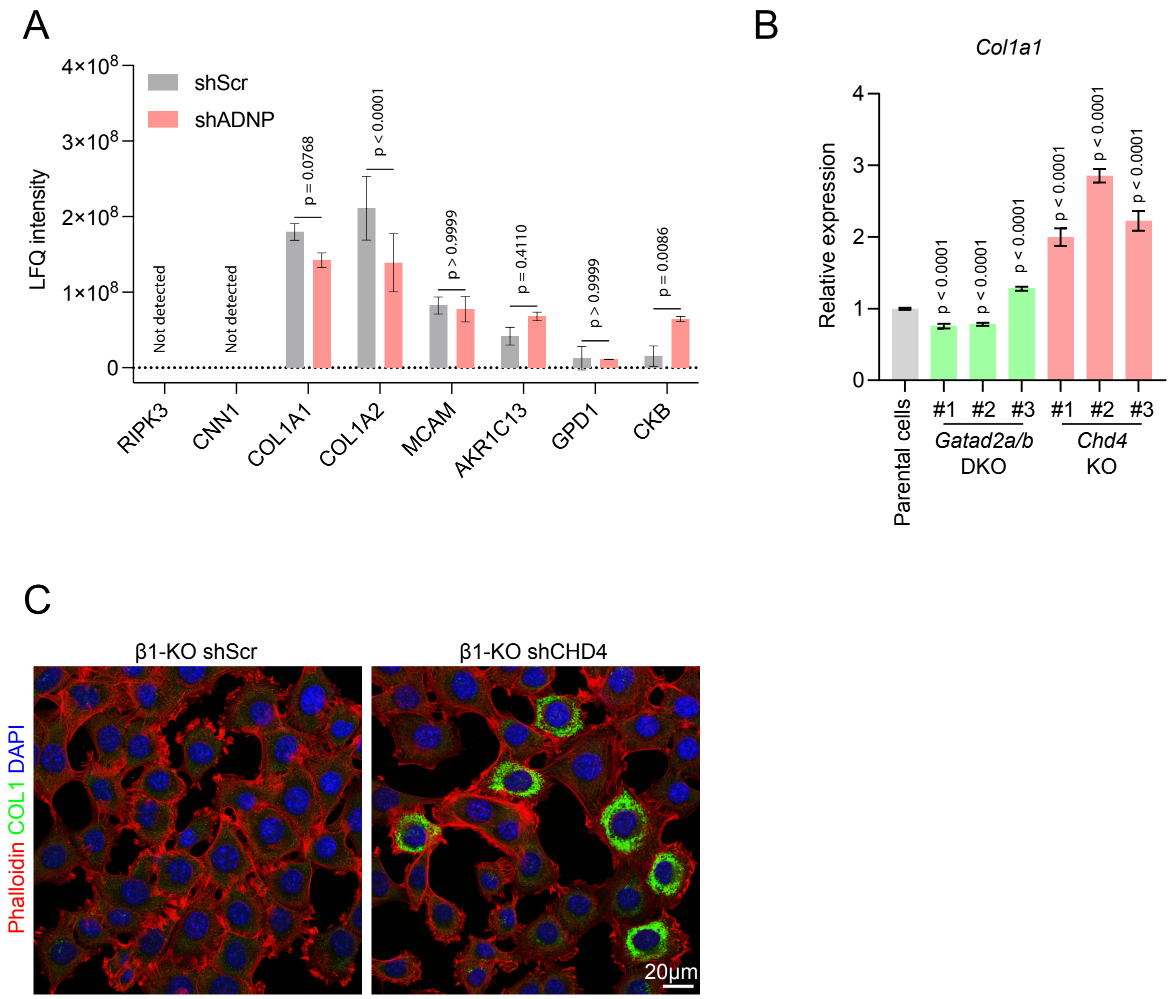

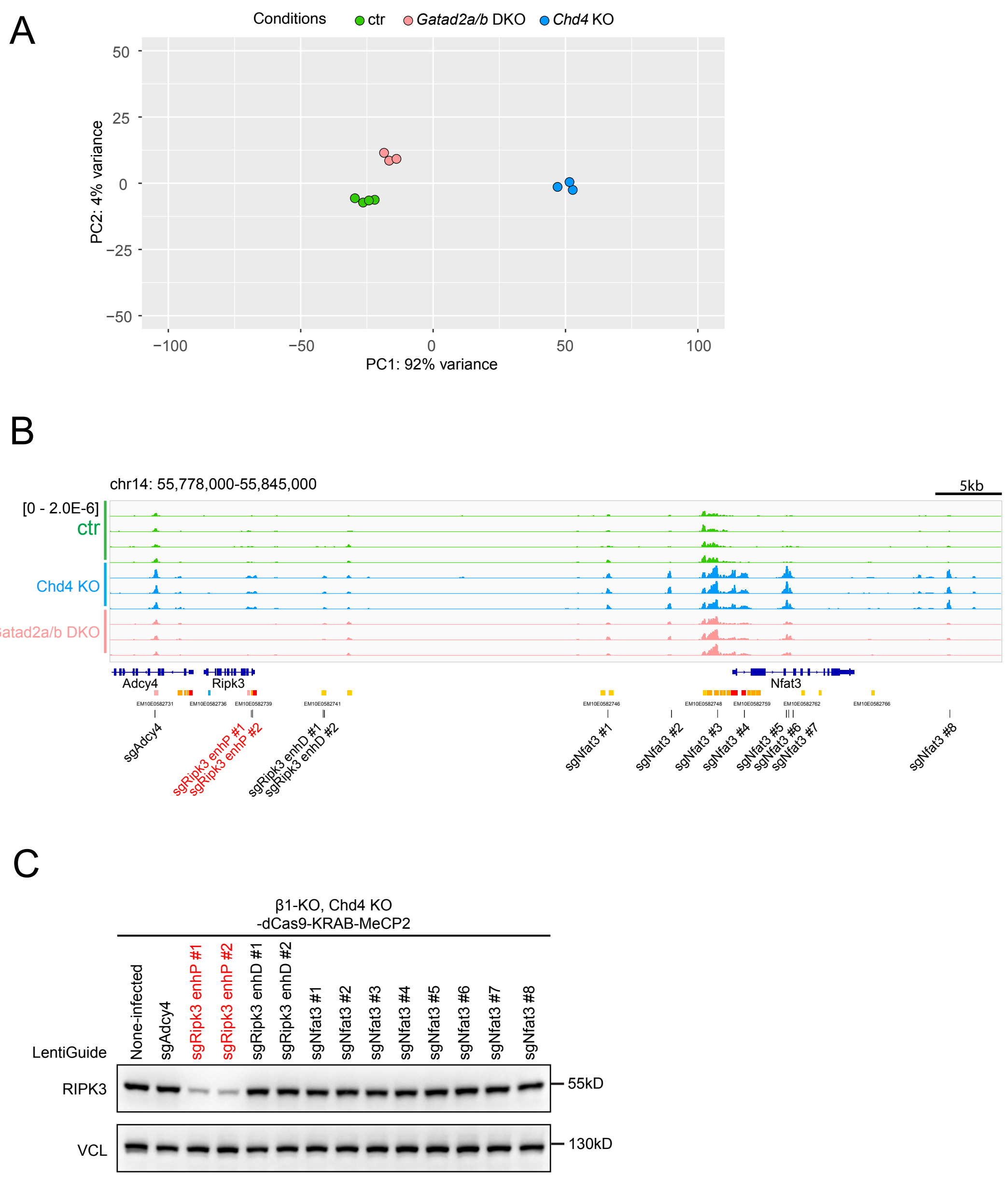
